# Selective control of prefrontal neural timescales by parietal cortex

**DOI:** 10.1101/2024.09.30.615928

**Authors:** Orhan Soyuhos, Marc Zirnsak, Rishidev Chaudhuri, Xiaomo Chen

## Abstract

Intrinsic neural timescales quantify how long spontaneous neuronal activity patterns persist, reflecting dynamics of endogenous fluctuations. We measured intrinsic timescales of frontal eye field (FEF) neurons and examined their changes during posterior parietal cortex (PPC) inactivation. We observed two distinct classes of FEF neurons based on their intrinsic timescales: short-timescale neurons (∼25 ms) or long-timescale neurons (∼100 ms). Short-timescale neurons showed stronger transient visual responses, suggesting their role in rapid visual processing, whereas long-timescale neurons exhibited pronounced sustained attentional modulation, suggesting their role in maintaining stimulus-driven attention. During PPC inactivation, intrinsic timescales increased in both neuron types, with a significantly larger effect in short-timescale neurons. In addition, PPC inactivation reduced attentional modulation, particularly in long-timescale neurons. Our findings provide the first causal evidence linking intrinsic local neural timescales to long-range inter-area communications. These findings also suggest the presence of at least two distinct network motifs that support different neuronal dynamics and functional computations within the FEF.

## Introduction

Intrinsic neural timescales measure the duration over which spontaneous neural activity remains temporally correlated with its prior states, independent of external stimuli or specific behavioral tasks (Cavanagh et al., 2020; Murray et al., 2014; Wolff et al., 2022). They define the temporal window in which previous neural activity can influence current neural dynamics, thereby shaping how information is processed and integrated within neural circuits (Hasson et al., 2015). These timescales exhibit systematic variations across brain regions, reflecting their functional specialization. For instance, along the visual hierarchy, early visual areas exhibit shorter intrinsic timescales, whereas higher-order areas demonstrate longer timescales (Chaudhuri et al., 2015; Murray et al., 2014). This gradient aligns with functional hierarchies in the brain, where shorter timescales are associated with regions involved in transient information processing, and longer timescales support sustained cognitive functions, such as working memory and decision-making (Cavanagh et al., 2016; Raut et al., 2020; Wasmuht et al., 2018; Trepka et al., 2024). Recent studies further suggest that intrinsic timescales also vary among individual neurons within a single brain region, corresponding to the functional heterogeneity at the neuronal level (Cavanagh et al., 2020; Zeraati et al., 2023; Spitmaan et al., 2020). Thus, this diversity may critically enable the brain’s dynamic balance between integrating and segregating information over multiple temporal scales (Cavanagh et al., 2020; Golesorkhi et al., 2021; Hasson et al., 2015; Wolff et al., 2022; Soltani et al., 2021).

In the primate brain, the frontal eye field (FEF) and the posterior parietal cortex (PPC) are key nodes in the frontoparietal attention network, with the PPC located earlier in the visual hierarchy (Buschman and Miller, 2007; Boshra and Kastner, 2022; Fiebelkorn and Kastner, 2020). Both areas are involved in stimulus-driven and goal-directed visual attention (Moore and Zirnsak, 2017; Xia et al., 2024). Our recent research demonstrated a causal role of the PPC in regulating stimulus-driven attentional representations in the FEF and attention-driven behavior (Chen et al., 2020). Anatomically, the FEF and oculomotor areas within the PPC, such as the lateral intraparietal area (LIP), exhibit extensive reciprocal connectivity (Lynch and Tian, 2006). These interactions between FEF and PPC are thought to play a crucial role in supporting various cognitive functions, including visual attention (Murray et al., 2017; Hart and Huk, 2020; Szczepanski et al., 2010; Fiebelkorn et al., 2018). Moreover, the FEF has direct anatomical connections with most visual cortical areas, both receiving input and sending feedback projections across the visual cortex (Schall et al., 1995; Stanton et al., 1995). Therefore, FEF is considered a proxy for prefrontal cortex communication with visual areas through these anatomical connections (Anderson et al., 2011). Consequently, intrinsic timescales within FEF likely play an important role in shaping attentional control across prefrontal and visual cortices.

The mechanisms underlying the variability in intrinsic neural timescales are thought to involve both inter- and intra-area interactions (Wang, 2020). Previous studies have primarily emphasized the role of local and regional factors within the brain, such as dendritic spine density in pyramidal neurons (Elston, 2000, 2003; Elston et al., 2011), the expression levels of NMDA and GABA receptor genes (Gao et al., 2020; Wang, 2008; Wong and Wang, 2006), and the extent of structural and functional connectivity (Hart and Huk, 2020; Fallon et al., 2020). Computational modeling studies have further shown that differences in intrinsic timescales across cortical regions can emerge naturally from variations in both local excitatory-inhibitory interactions and long-range connectivity patterns (Chaudhuri et al., 2014, 2015; Demirtaş et al., 2019; Wang, 2020). These findings suggest that intrinsic timescales arise from a combination of diverse excitatory and inhibitory connection strengths within regions and long-range connectivity patterns, indicating that neither factor alone is sufficient to predict a region’s timescales (Chaudhuri et al., 2014, 2015; Litwin-Kumar and Doiron, 2012). However, despite the modeling work, a direct causal link between inter-area communication and intrinsic neural timescales remains untested. Specifically, it is entirely unknown whether the intrinsic timescales of FEF neurons depend on PPC input.

In this study, we examined the diversity of intrinsic neural timescales among FEF neurons, assessed their relationship to neuronal functional specialization during visual processing and attention tasks, and investigated their dependence on PPC inputs. We found a bimodal distribution of intrinsic timescales, identifying distinct FEF neuronal groups with either fast (short-*τ*) or slow (long-*τ*) timescales. Moreover, these timescales measured during the baseline period correlated with the neurons’ functional properties during the task: short-*τ* neurons were more involved in processing transient visual input, while long-*τ* neurons exhibited stronger sustained stimulus-driven attentional modulation. Finally, PPC inactivation selectively increased the intrinsic timescales of short-*τ* neurons while having a larger impact on attentional modulation of long-*τ* neurons. These findings provide direct causal evidence that PPC inputs regulate intrinsic dynamics and attentional modulation within FEF, suggesting a distributed computation mechanism underlying the frontoparietal attention network.

## Results

We recorded spiking activity within the FEFs in two behaving monkeys (J and Q) using multi-channel microelectrodes (Fig. 1A; see Methods). In each trial, neuronal responses were measured during presentation of different visual stimuli, including single visual probes (“Single”) and popout arrays containing unique stimuli (“Popout”) (Fig. 1B). Additionally, to assess the impact of posterior parietal cortex (PPC) inputs on FEF activity, we reversibly inactivated PPC using cryoloops chronically implanted within the intraparietal sulcus (IPS). We then compared FEF spiking responses between PPC inactivation (“Inactivation”) and control (“Control”) conditions. In total, we recorded 400 single- and multi-unit responses from 192 FEF recording sites across 11 experimental sessions.

**Fig. 1.**
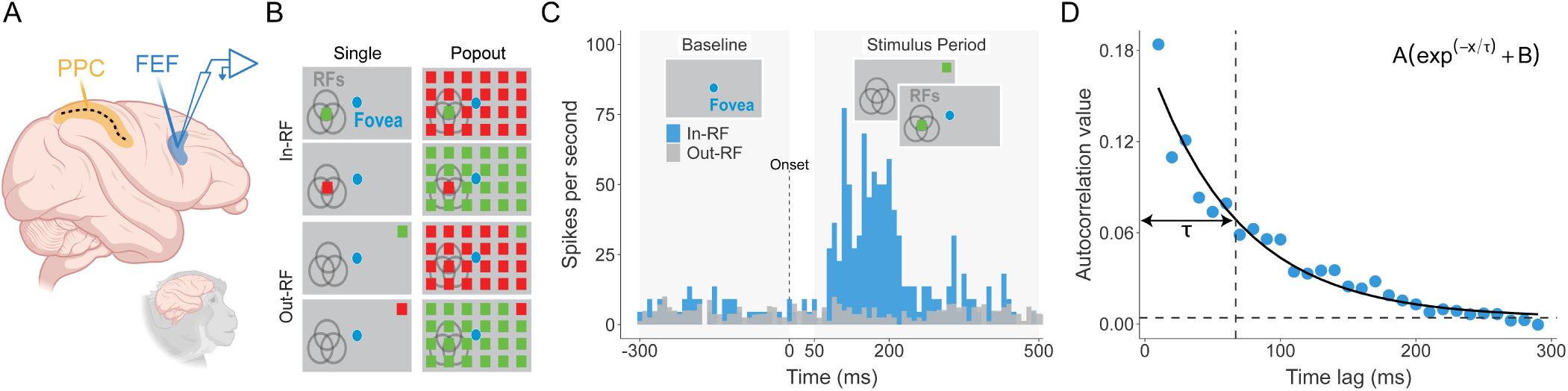
Experimental setup and measurement of intrinsic neural timescales. (A) Schematic illustration of the experimental setup. Neuronal activity in the frontal eye field (FEF) was recorded using a linear electrode array under two conditions: control and during cryo- inactivation of the posterior parietal cortex (PPC). (B) Visual stimulus conditions included either a single colored stimulus presented alone (“Single”) or the same stimulus embedded among identically colored distractors (“Popout”). (C) Example neuronal response illustrating mean firing rate to a single stimulus presented either inside (blue trace) or outside (gray trace) the neuron’s receptive field (RF). (D) Intrinsic neural timescales were measured using spike-count autocorrelation computed from neuronal activity during the baseline period. Blue dots indicate autocorrelation coefficients at successive time lags. Timescales were derived by fitting an exponential decay function (solid curve) to the autocorrelation data, where the resulting decay constant (*τ*; vertical dashed line) reflects the intrinsic neural timescale, quantifying how quickly neuronal activity becomes decorrelated over time. The horizontal dashed line indicates the offset of the exponential fit.

### Two distinct neural timescales in FEF neurons

To characterize the diversity of intrinsic neuronal dynamics within the FEF, we first analyzed neural timescales in two monkeys under the control condition (no PPC inactivation). Intrinsic neural timescales were quantified by calculating the spike-count autocorrelation for each neuron based on spontaneous activity in the baseline epoch before stimulus presentation (Fig. 1C). Autocorrelation functions were fit using a single exponential decay model to estimate intrinsic neural timescales (Fig. 1D) (Murray et al., 2014). R-squared (*R*^2^) values were used to quantify the goodness of fit between our model and the observed data (Fig. S1A). The exponential function was closely aligned with the autocorrelation profiles of each neuron (mean *R*^2^ = 0.78; *R*^2^ for Monkey J = 0.73; *R*^2^ for Monkey Q = 0.83). Neurons with poor fits (*R*^2^ ¡ 0.3, n = 20) were excluded, leaving 380 neurons for analysis. A relatively long-*τ* indicated that baseline neural activity was sustained over extended periods with a stable firing pattern, whereas a short-*τ* signified more transient activity characterized by a dynamic baseline firing pattern.

The distribution of intrinsic timescales among FEF neurons exhibited clear bimodality (Fig. 2A), confirmed by several statistical tests (Table S1). This bimodality was observed across both monkeys combined (Excess Mass Test: *p* = 2 × 10*^−^*^3^) and within each monkey individually (Excess Mass Test: Monkey Q, *p <* 2.2 × 10*^−^*^16^; Monkey J, *p* = 2 × 10*^−^*^2^) (Ameijeiras-Alonso et al., 2019). Neurons were divided into short-*τ* and long-*τ* populations using the global minimum (60.95 ms) from the kernel density estimate of the combined distribution. These global minima were consistent across individual monkeys (Monkey J = 60.51 ms; Monkey Q = 59.33 ms) (Fig. 2B). Consequently, neurons in the short-*τ* group exhibited a rapid decay rate (N = 168, mean = 27.34 ms), whereas neurons in the long-*τ* group showed a slower decay rate in their auto-correlation values during the baseline period (N = 212, mean = 102.64 ms) (Fig. 2C). Similar distributions were observed for both Monkey J and Monkey Q (short-*τ* neurons: Monkey J = 28.17 ms; Monkey Q = 26.71 ms; long-*τ* neurons: Monkey J = 104.78 ms; Monkey Q = 100.73 ms). In contrast, firing rate distributions exhibited a unimodal pattern (Excess Mass Test: Monkey Q, *p* = 0.53; Monkey J, *p* = 0.78) (Fig. S5B). Together, these results suggest the existence of at least two distinct neural circuit motifs within the FEF characterized by different temporal dynamics.

**Fig. 2.**
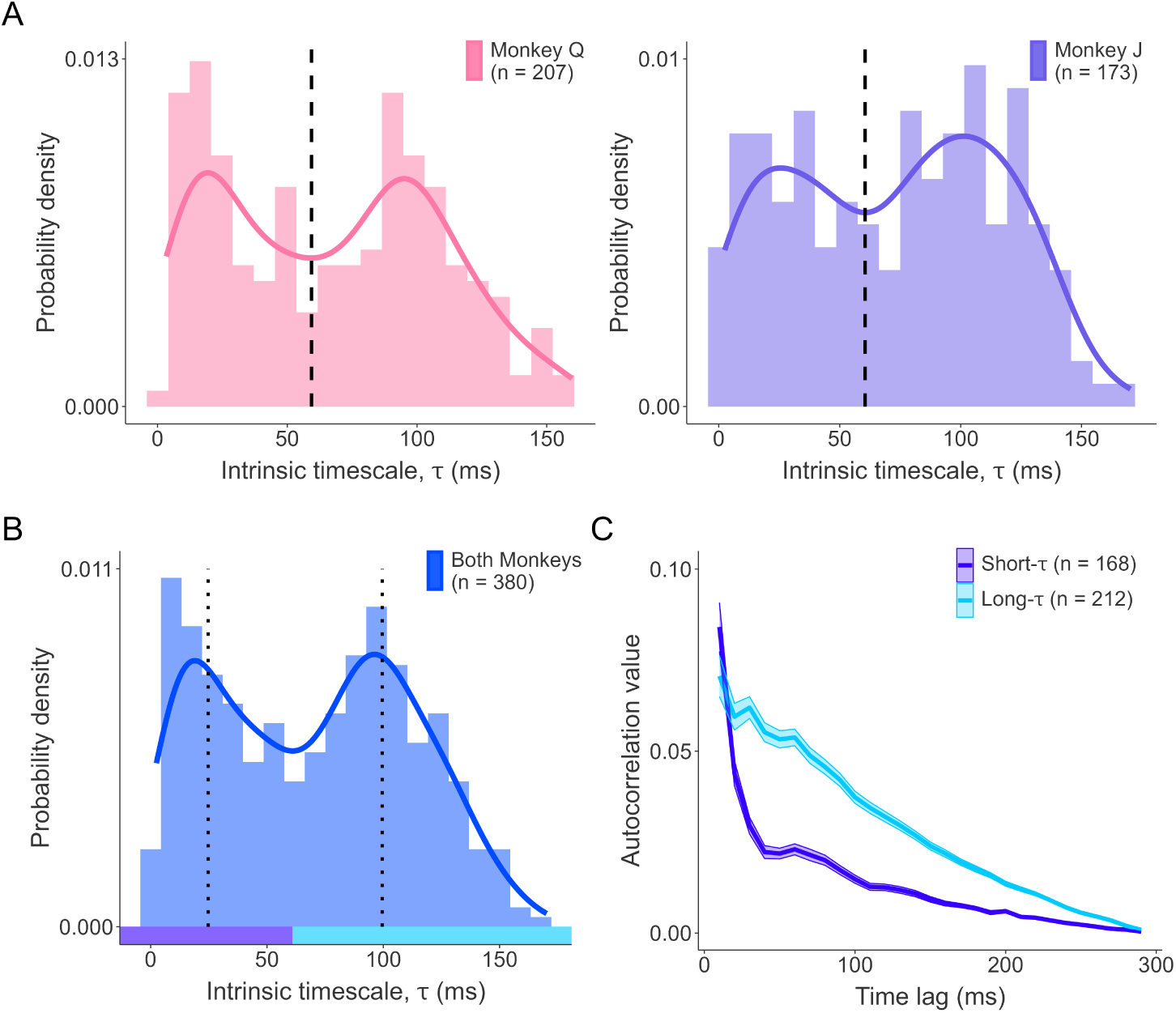
Bimodal distribution of intrinsic neural timescales in FEF neurons. (A) Histograms and kernel density estimates of intrinsic neural timescales (*τ*) for monkey Q (left) and monkey J (right). Solid lines represent the kernel density estimates. Dashed lines mark the global minima separating neurons into short-timescale (short-*τ*) and long-timescale (long-*τ*) groups. (B) Histogram and kernel density estimate of *τ* values for both monkeys combined. Dotted lines denote the median *τ* for each group. (C) Mean autocorrelation profiles computed for short-*τ* (purple) and long-*τ* (blue) neurons. Shaded areas indicate ± the standard error of the mean (SEM).

### Functional relevance of FEF intrinsic neural timescales

Next, we asked whether the intrinsic neural timescales measured during the task-free baseline period correlate with the neurons’ functional properties during the task. We used single and popout indices to quantify neuronal visual responses and attentional modulation. Specifically, the single index was calculated by comparing the activity evoked by a single stimulus presented inside versus outside the receptive field (RF). This index captures the spatial selectivity of visual responses, reflecting the ability of neurons to distinguish stimuli within their RFs. Similarly, a popout index was calculated by comparing neuronal responses to unique stimuli inside versus outside the RF. This index quantifies stimulus-driven attentional modulation, reflecting the ability of neurons to distinguish visually salient stimuli. To further characterize the temporal dynamics of these responses, we divided the neuronal activity into transient (50–200 ms) and sustained (200–500 ms) periods following stimulus onset. This temporal segmentation allowed us to calculate transient and sustained versions of both single and popout indices, enabling us to distinguish rapid, short-lived neural responses to stimulus onset from long-term changes in neural activity associated with sustained modulation (see Methods).

To quantify the functional relevance of FEF neurons’ intrinsic timescales, we used multiple linear regression analysis to identify relationships between these timescales and both single and popout indices during transient and sustained epochs (Fig. 3A). Here, multiple linear regression analysis allowed us to assess the correlation between intrinsic neural timescales and each predictor, while controlling for the influence of other predictors. For visual responses, our results revealed a significant negative association between timescales and transient single indices (Permutation test: *p* = 2 × 10*^−^*^4^, *β* = −73.98). That is, neurons with fast intrinsic timescales showed stronger responses to single visual stimuli within their receptive field than neurons with slower intrinsic timescales. For attentional modulation, there was a significant negative relationship between timescales and transient popout indices (Permutation test: *p* = 2.05 × 10*^−^*^2^, *β* = −53.98), as well as a significant positive relationship with sustained popout indices (Permutation test: *p* = 9 × 10*^−^*^4^, *β* = 77.18). In other words, neurons with slower intrinsic timescales exhibited stronger sustained attentional modulation, while neurons with faster timescales showed greater transient attentional modulation. These findings demonstrate a robust correlation between individual FEF neurons’ intrinsic timescales and both their visual responses and attentional modulation.

**Fig. 3.**
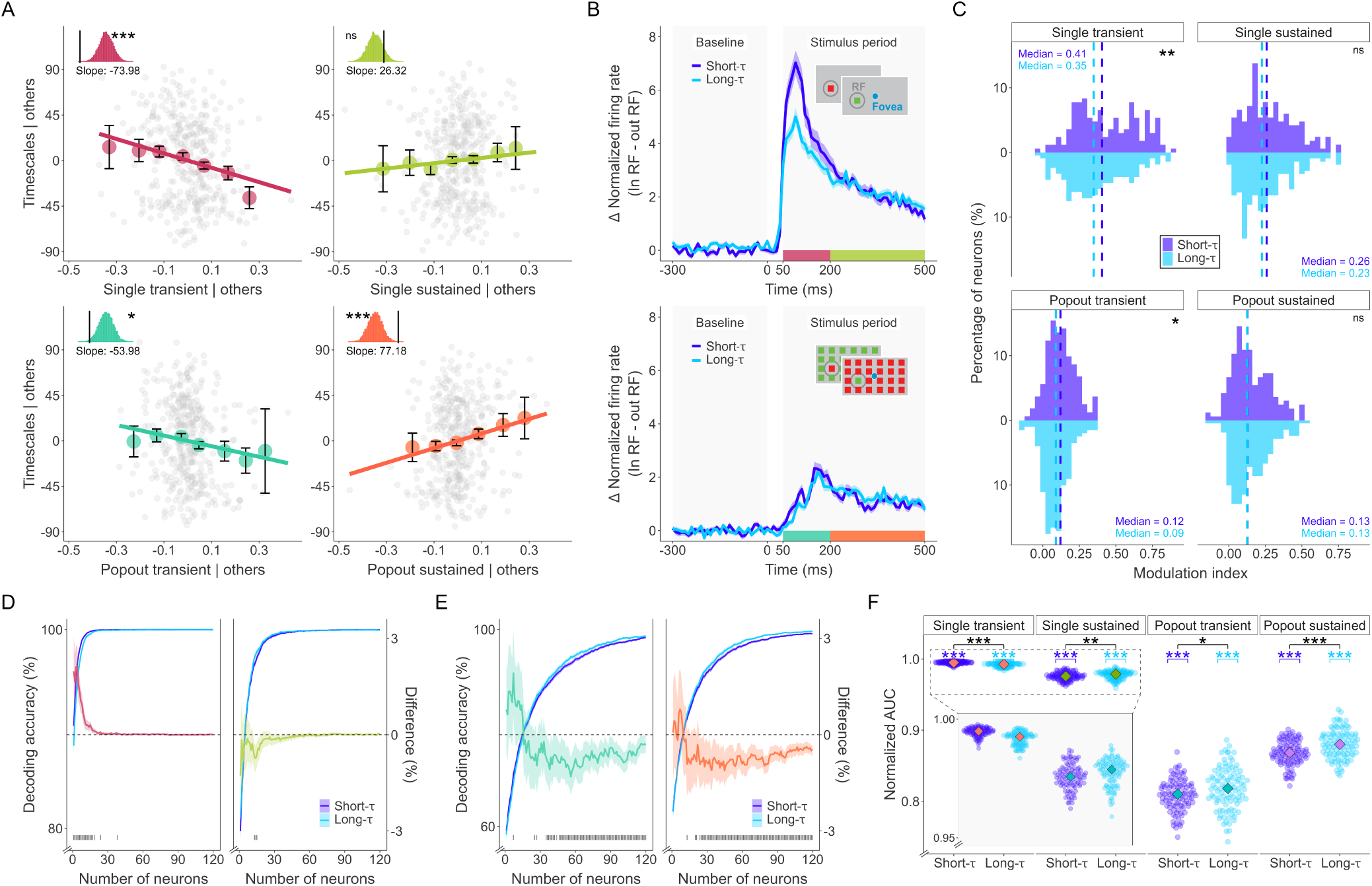
Relationship between intrinsic neural timescales and functional properties of FEF neurons. (A) Added-variable plots illustrating the relationship between intrinsic neural timescales (*τ*) and single (top) or popout (bottom) indices during transient (left) and sustained (right) response periods. Insets indicate significance levels determined by nonparametric permutation tests. (B) Differences in normalized neuronal responses evoked by single (top) and popout stimuli (bottom) presented inside (In-RF) versus outside the receptive fields (Out-RF), shown separately for short-*τ* (purple) and long-*τ* (blue) neurons. Shaded areas indicate ± SEM. (C) Distributions of single (top) and popout indices (bottom) during transient (left) and sustained (right) periods, shown separately for short-*τ* (purple) and long-*τ* (blue) neuronal groups. Dashed lines indicate median values. (D) Population decoding performance for discriminating single stimuli presented inside versus outside RFs as a function of population size. Results shown separately for transient (left) and sustained (right) periods and for short-*τ* (purple) and long-*τ* (blue) neurons. Secondary axes (red and green lines) indicate performance differences (long-*τ* minus short-*τ*) as a function of population size, with shaded ribbons representing 95% confidence intervals (CIs). Gray vertical lines mark significant differences. The black dashed line represents chance-level decoding performance. (E) Population decoding performance for discriminating popout stimuli presented inside versus outside RFs, formatted similarly to panel D. Performance differences between long-*τ* and short-*τ* groups are shown by turquoise and orange lines. (F) Summary comparison of normalized area under the curve (AUC) values for population decoding performance between short-*τ* (purple) and long-*τ* (blue) neuron groups. Diamonds indicate median values; the inset shows single-stimulus decoding results at an expanded scale. (*, *p <* 0.05; **, *p <* 0.01; ***, *p <* 0.001; ns = not significant).

Given the clear functional relevance of FEF intrinsic timescales, we next asked whether visual responses and attentional modulation differ between two distinct groups of FEF neurons, defined by their intrinsic timescales during the task-free epoch (Fig. 3B). To answer this question, we first compared the single and popout indices between short-*τ* and long-*τ* neurons. The distributions for transient single indices (Cramer–von Mises test: *p <* 3 × 10*^−^*^3^) and transient popout indices (Cramer–von Mises test: *p <* 1.3 × 10*^−^*^2^) were significantly greater for short-*τ* neurons compared to long-*τ* neurons. In contrast, sustained indices were not significantly different for either single (Cramer–von Mises test: *p* = 0.16) or popout (Cramer–von Mises test: *p* = 0.96) at the individual level (Fig. 3C). Additionally, the visual onsets for the two groups did not significantly differ from each other for single (Cramer–von Mises test: *p* = 0.19) or popout stimuli (Cramer–von Mises test: *p* = 0.08) (Fig. S6A). These findings align with the results from the regression analyses, demonstrating that neurons with shorter timescales exhibit stronger, but not faster, transient visual responses and attentional modulation.

Next, we compared the amount of visual and attentional information that short-*τ* and long-*τ* neurons had at the population level and performed pseudo-population decoding analysis to predict the location of a single stimulus inside or outside the RFs (single-decoding) and the location of a popout stimulus inside or outside the RFs (popout-decoding) (see Methods). We generated neuron-dropping curves (NDCs) (Lebedev, 2014; Pettine et al., 2019; Wessberg et al., 2000) using the performance of neuronal subsets obtained from either short-*τ* or long-*τ* neurons to examine decoding accuracies across varying population sizes. Both groups had high decoding accuracy, consistently above chance level across all population sizes, but differed in their performance profiles (Fig. 3D–E).

To quantify their differences in decoding performance, we calculated the area under the curve (AUC) exceeding the chance level for each group to assess population decoding performance (Fig. 3F; complementary findings from the saturation fitting analysis are also available in Fig. S3). Both single- and popout-decoding performances differed significantly between short-*τ* and long-*τ* neurons across both transient and sustained periods. Short-*τ* neurons exhibited higher AUC values for single-decoding during the transient period (Wilcoxon rank-sum test: *p* = 2.83 × 10*^−^*^13^, effect size = 0.60; median = 0.995 for short-*τ* vs. 0.993 for long-*τ*). In comparison, long-*τ* neurons exhibited better single-decoding performance during the sustained period (Wilcoxon rank-sum test: *p* = 8.41 × 10*^−^*^3^, effect size = 0.22; median = 0.979 for long-*τ* vs. 0.976 for short-*τ*). Additionally, long-*τ* neurons had greater popout-decoding performance during both the transient (Wilcoxon rank-sum test: *p* = 1.49 × 10*^−^*^2^, effect size = 0.20; median = 0.818 for long-*τ* vs. 0.810 for short-*τ*) and sustained periods (Wilcoxon rank-sum test: *p* = 1.98 × 10*^−^*^5^, effect size = 0.35; median = 0.880 for long-*τ* vs. 0.868 for short-*τ*). The differences between the groups were most pronounced for single-decoding in the transient period (effect size = 0.60) and popout-decoding in the sustained period (effect size = 0.35). These results highlight the complementary roles of short-*τ* and long-*τ* neurons in visual and attentional modulation, with short-*τ* neurons specialized for short-lived, transient visual responses and long-*τ* neurons for sustained attentional modulation.

In summary, intrinsic neural timescales in the FEF are closely linked to visual response and attentional modulation at both the individual and population levels, with short-*τ* neurons excelling in transient visual processing and long-*τ* neurons playing a greater role in sustained attentional modulation.

### Selective increase of FEF intrinsic neural timescales during PPC inactivation

We then investigated the effects of PPC inactivation on the FEF intrinsic timescales to understand how PPC regulates local neuronal dynamics within the FEF. PPC inactivation was achieved using cryoloops implanted in the IPS, which suppressed neuronal activity by lowering the temperature of the cortical tissue. The effectiveness of PPC inactivation was confirmed by robust behavioral effects in both the free viewing task and the double-target choice task during inactivation (Chen et al., 2020). Similar to the control condition, the exponential function showed a good fit to each neuron’s autocorrelation profiles during inactivation (mean *R*^2^ = 0.80; *R*^2^ for Monkey J = 0.75; *R*^2^ for Monkey Q = 0.84; Fig. S1B). Fig. 4A shows six representative neurons, illustrating the changes in their timescales during PPC inactivation. These neurons exhibited an increase in their intrinsic timescales, reflecting slower dynamics and a more stable firing pattern during the baseline period. Consistent with this, we found a significant increase in intrinsic timescales across the entire population of FEF neurons following PPC inactivation (Wilcoxon signed-rank test: *p* = 5.09 × 10*^−^*^13^, effect size = 0.432; mean = 84.43 ms, median = 94.92 ms for inactivation vs. mean = 69.89 ms, median = 73.14 ms for control) (Fig. 4B). This increase in intrinsic neural timescales was evident in both monkeys (Wilcoxon signed-rank test: *p* = 3.02 × 10*^−^*^2^, effect size = 0.19 for Monkey J; *p* = 4 × 10*^−^*^15^, effect size = 0.63 for Monkey Q). Thus, PPC inactivation robustly slowed neural dynamics across FEF neurons.

**Fig. 4.**
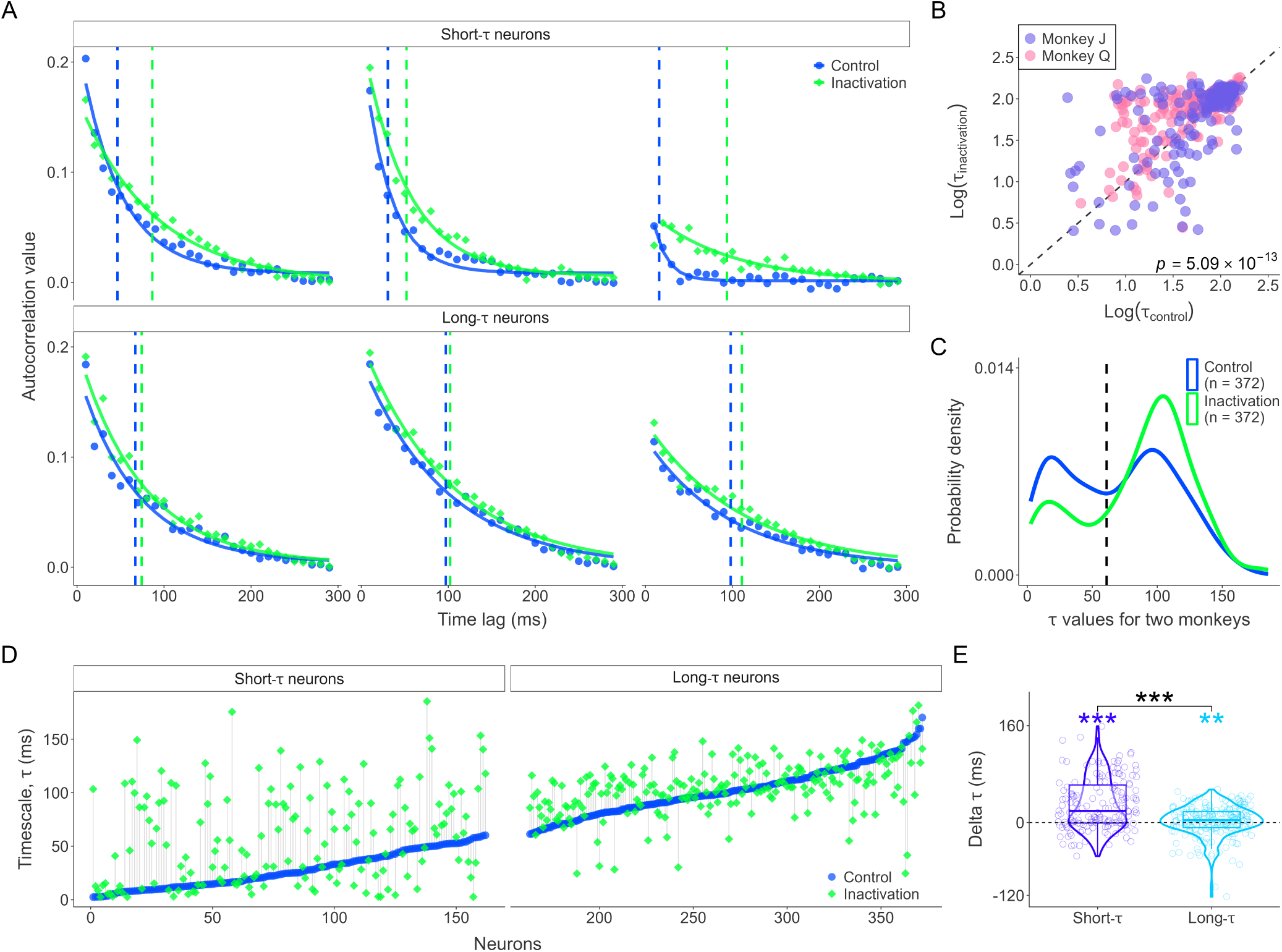
PPC inactivation selectively alters intrinsic neural timescales of FEF neurons. (A) Example autocorrelation profiles from representative short-*τ* (top) and long-*τ* (bottom) neurons during control (blue) and posterior parietal cortex (PPC) inactivation (green) conditions. Dashed vertical lines indicate intrinsic neural timescales (*τ*). (B) Scatter plot comparing log-transformed *τ* values between control and PPC inactivation conditions across both monkeys (Monkey J, purple; Monkey Q, pink). (C) Probability density estimates illustrating the distribution of *τ* under control (blue) and PPC inactivation (green) conditions. (D) Individual neurons sorted by their *τ* values under control condition, shown separately for short-*τ* (left) and long-*τ* (right) neuronal groups. Blue and green markers represent control and PPC inactivation conditions, respectively. (E) Violin plots depicting changes in *τ* values (Δ*τ*, inactivation minus control) between short-*τ* and long-*τ* neuron groups following PPC inactivation. (*, *p <* 0.05; **, *p <* 0.01; ***, *p <* 0.001; ns = not significant).

We further explored the changes in intrinsic timescales separately for the two distinct populations in FEF: short-*τ* and long-*τ* neurons. Similar to control conditions, we found that the probability density of FEF timescales exhibited a bimodal distribution during PPC inactivation (Excess Mass Test: *p <* 2.2 × 10*^−^*^16^). However, PPC inactivation led to a substantial increase (29.52%) in the number of long-*τ* neurons (*N*_Control_ = 210; *N*_Inactivation_ = 272) and a corresponding decrease (−38.27%) in the number of short-*τ* neurons (*N*_Control_ = 162; *N*_Inactivation_ = 100). This redistribution highlights a shift in the population dynamics toward slower timescales (Fig. 4C).

We next examined the magnitude of timescale changes separately for short-*τ* and long-*τ* neurons (Fig. 4D). We found that there was a significant increase in timescales for both short-*τ* neurons (Wilcoxon signed-rank test: *p* = 1.02 × 10*^−^*^13^, effect size = 0.67) and long-*τ* neurons (Wilcoxon signed-rank test: *p* = 7.79 × 10*^−^*^3^, effect size = 0.21). However, the changes observed in short-*τ* neurons were considerably larger than those of long-*τ* neurons. On average, short-*τ* neurons exhibited a 15-fold greater increase in timescales compared to long-*τ* neurons (Wilcoxon rank-sum test: *p* = 8.64 × 10*^−^*^9^, effect size = 0.35; short-*τ* neurons: Δ*τ* mean = 30 ms, Δ*τ* median = 19 ms; long-*τ* neurons: Δ*τ* mean = 2 ms, Δ*τ* median = 4 ms) (Fig. 4E). This effect was also observed individually in each monkey (Monkey J: *p* = 3.65 × 10*^−^*^2^, effect size = 0.19; Monkey Q: *p* = 4.10 × 10*^−^*^9^, effect size = 0.48). Notably, long-*τ* neurons in Monkey J did not show a significant change in their timescales (*p* = 0.62). Additionally, the changes in intrinsic neural timescales were not correlated with the changes in baseline firing rate (Spearman’s rank correlation: *r* = 0.09, *p* = 0.07) (Fig. S5A). Although baseline firing rates slightly increased following PPC inactivation (Wilcoxon signed-rank test: *p* = 5.94 × 10*^−^*^3^, Δ firing rate = 0.93 Hz; Fig. S5B), these changes were similar between short-*τ* and long-*τ* neurons (Wilcoxon rank-sum test: *p* = 0.33) (Fig. S5C). These results indicate a selective dependence of FEF neural dynamics on PPC input.

### PPC inactivation selectively weakens the stimulus-driven attentional modulation in FEF neurons

Finally, we assessed the effects of PPC inactivation on stimulus-driven attention modulation. We first analyzed the relationship between neural timescales and the functional properties of FEF neurons during PPC inactivation. Similar to the control condition, we employed multiple linear regression analysis to assess the correlation between intrinsic neural timescales and both single and popout indices during inactivation conditions (Fig. 5A and S4). Our results revealed that the correlation between intrinsic timescales and attentional modulation relies on the input from PPC during both transient and sustained periods. Specifically, the significant negative association between transient popout and intrinsic timescales in the control condition (Permutation test: *p* = 2.05 × 10*^−^*^2^, *β* = −53.98) was no longer significant during PPC inactivation (Permutation test: *p* = 0.63, *β* = −12.45). Additionally, the positive association between sustained popout indices and intrinsic timescales under control conditions (Permutation test: *p* = 9 × 10*^−^*^4^, *β* = 77.18) became nonsignificant during inactivation (Permutation test: *p* = 0.21, *β* = −32.12). For visual modulation, PPC inactivation reduced but did not eliminate the significant negative correlation between intrinsic timescales and transient single indices (Permutation test: *p* = 2 × 10*^−^*^4^, *β* = −73.98 under control; *p* = 2.53 × 10*^−^*^2^, *β* = −44.83 after inactivation). No significant relationship was found between intrinsic timescales and sustained single indices in either condition (Permutation test: *p* = 0.24, *β* = 26.32 under control; *p* = 0.54, *β* = 13.77 after inactivation). Thus, PPC inactivation exerted a larger impact on the correlation between neural timescales and attentional modulation compared to visual response.

**Fig. 5.**
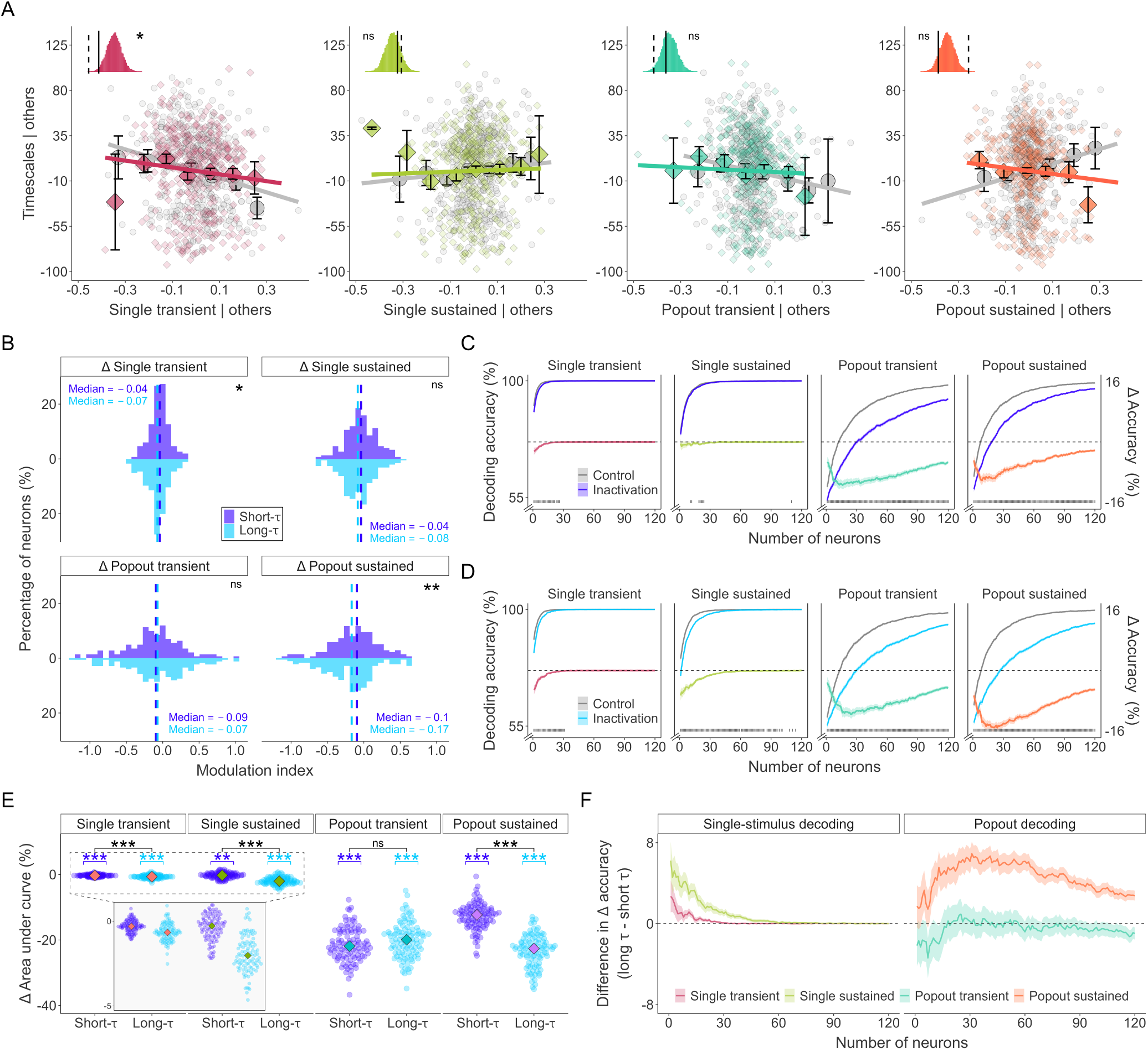
Effects of PPC inactivation on visual responses and attentional modulation in FEF neurons. (A) Added-variable plots illustrating correlations between intrinsic neural timescales (*τ*) and single/popout indices before (gray circles) and after posterior parietal cortex (PPC) inactivation (colored diamonds). Insets depict slopes fitted to data before (solid lines) and after (dashed lines) PPC inactivation compared to null distributions obtained from permutation tests. (B) Change (Δ) in single and popout indices (inactivation minus control) for short-*τ* (purple) and long-*τ* (blue) neuronal groups following PPC inactivation. Dashed horizontal lines indicate median Δ values. (C) Population decoding performance for discriminating single (left two plots) and popout (right two plots) stimuli presented inside versus outside receptive fields (RF) as a function of neuronal population size in short-*τ* neurons. Performance is shown separately for transient and sustained response periods during control (gray) and PPC inactivation (purple). Colored lines on secondary axes show differences in decoding performance (inactivation minus control; red: single transient, green: single sustained, turquoise: popout transient, orange: popout sustained). Gray vertical bars indicate statistically significant differences. (D) Population decoding performance for single and popout stimuli, formatted as in (C), for long-*τ* neurons under control (gray) and PPC inactivation (blue) conditions. (E) Comparison of changes in decoding performance (Δ area under the curve [AUC], inactivation minus control) between short-*τ* (purple) and long-*τ* (blue) neuronal groups. Diamonds indicate median values; the inset magnifies changes in decoding performance for a single stimulus. (F) Differences in decoding performance changes between long-*τ* and short-*τ* neurons following PPC inactivation. Positive values indicate a greater reduction in decoding performance in long-*τ* neurons, whereas negative values indicate a greater reduction in short-*τ* neurons. (*, *p <* 0.05; **, *p <* 0.01; ***, *p <* 0.001; ns = not significant).

We next assessed how PPC inactivation influenced visual responses and attentional modulation in individual FEF neurons, and examined whether these effects differed between short-*τ* and long-*τ* neurons. We quantified this impact using the changes in single and popout indices for each neuron. Consistent with the linear regression analysis and our previous findings (Chen et al., 2020), PPC inactivation had a significantly larger impact on popout indices than single indices during the sustained period (Wilcoxon rank-sum test: *p* = 2.17 × 10*^−^*^4^, effect size = 0.22; Δsingle index: mean = –0.04, median = –0.05; Δpopout index: mean = –0.13, median = –0.13).

We then examined the effects on short-*τ* and long-*τ* neurons separately. PPC inactivation had a greater effect on long-*τ* neurons than short-*τ* neurons for both transient single (Cramer–von Mises test: *p* = 1.70 × 10*^−^*^2^) and sustained popout indices (Cramer–von Mises test: *p <* 10*^−^*^2^) (Fig. 5B). Additionally, the visual onsets of both groups were significantly delayed following PPC inactivation for both single (Cramer–von Mises test: *p* = 7 × 10*^−^*^3^ for short-*τ*, Δvisual onset: mean = 11 ms, median = 10 ms; *p <* 10*^−^*^3^ for long-*τ*, Δvisual onset: mean = 23 ms, median = 10 ms) and popout stimuli (Cramer–von Mises test: *p <* 10*^−^*^3^ for short-*τ*, Δvisual onset: mean = 40 ms, median = 30 ms; *p* = 1.3 × 10*^−^*^2^ for long-*τ*, Δvisual onset: mean = 22 ms, median = 20 ms) (Fig. S6B–C).

We further examined the impact of PPC inactivation at the population level using the neuron-dropping curves (Fig. 5C–D). Consistent with individual neurons, PPC inactivation had a larger impact on attentional modulation than visual response and a larger impact on long-*τ* neurons compared to short-*τ* neurons (Fig. 5E). Specifically, for single-decoding performance, both short-*τ* and long-*τ* neurons showed slight but significant decreases in AUC above the chance level during the transient period. Short-*τ* neurons decreased by 0.31% (Wilcoxon signed-rank test: *p* = 4.64 × 10*^−^*^12^) and long-*τ* neurons decreased by 0.67% (Wilcoxon signed-rank test: *p* = 8.46 × 10*^−^*^16^), with long-*τ* neurons exhibiting a larger reduction (Wilcoxon rank-sum test: *p* = 5.94 × 10*^−^*^8^). During the sustained period, the decreases were 0.31% for short-*τ* (Wilcoxon signed-rank test: *p* = 1.12 × 10*^−^*^3^) and 1.97% for long-*τ* (Wilcoxon signed-rank test: *p* = 9.46 × 10*^−^*^18^), again showing a more pronounced decline in long-*τ* neurons (Wilcoxon rank-sum test: *p* = 1.01 × 10*^−^*^20^). In contrast, popout-decoding performance exhibited more substantial declines. Short-*τ* neurons experienced a 21.50% decrease in transient popout-decoding performance (Wilcoxon signed-rank test: *p* = 3.96 × 10*^−^*^18^) and a 12.84% reduction in sustained popout-decoding performance (Wilcoxon signed-rank test: *p* = 3.96 × 10*^−^*^18^), while long-*τ* neurons showed a 20.07% drop in transient popout-decoding performance (Wilcoxon signed-rank test: *p* = 3.96 × 10*^−^*^18^) and a 23.28% decrease in sustained popout-decoding performance (Wilcoxon signed-rank test: *p* = 3.96 × 10*^−^*^18^). Although both groups exhibited reductions, long-*τ* neurons demonstrated significantly greater decreases for popout-decoding performance in the sustained period (Wilcoxon rank-sum test: *p* = 1.23 × 10*^−^*^28^) (Fig. 5F).

In summary, PPC inactivation disrupted the relationship between intrinsic neural timescales and stimulus-driven attentional modulation, eliminating significant correlations with both transient and sustained popout indices. At both the individual and population levels, PPC inactivation predominantly impaired attentional modulation in long-*τ* neurons, particularly during the sustained attention period.

## Discussion

We found that intrinsic neural timescales within the FEF exhibit a bimodal distribution, revealing two functionally distinct neuronal populations of FEF neurons with fast (short-*τ*) and slow (long-*τ*) dynamics. These intrinsic timescales, measured during the baseline fixation period, correlated with neuronal functional properties. Specifically, short-*τ* neurons showed stronger transient responses to single visual stimuli, suggesting their important role in rapid visual processing, while long-*τ* neurons exhibited stronger sustained responses to popout arrays, suggesting their role in maintaining stimulus-driven attentional modulation over time. Importantly, PPC inactivation selectively influenced these populations by predominantly increasing intrinsic timescales in short-*τ* neurons and substantially reducing attentional modulation in long-*τ* neurons. These findings demonstrate that PPC inputs selectively shape intrinsic neuronal dynamics and functional properties within the FEF, supporting distributed computations across the frontoparietal attention network.

Our findings provide clear empirical evidence for two distinct neuronal classes in the FEF, characterized by their intrinsic neural timescales. This distinction is supported by consistent bimodal distributions observed across individual monkeys and by the selective effects of PPC inactivation. Importantly, these neuronal classes differed primarily in their temporal correlation rather than their initial response latencies, which remained comparable between groups (Fig. S6A). Short-*τ* neurons with shorter temporal correlation windows likely facilitate rapid updating of visual information, allowing them to segregate distinct events in dynamic environments (Chaudhuri et al., 2015; Murray et al., 2014). In contrast, long-*τ* neurons with their prolonged activity likely support stable integration across time and visual space, maintaining sustained attentional prioritization (Cavanagh et al., 2020; Wasmuht et al., 2018). Computationally, this heterogeneity may underpin the functional flexibility of the FEF (Cavanagh et al., 2020; Soltani et al., 2021), enabling it to effectively support diverse cognitive functions, including visually guided saccades (Bruce and Goldberg, 1985; Dias and Segraves, 1999), spatial attention (Hü er et al., 2024; Moore and Fallah, 2001, 2004; Thompson et al., 2005), decision-making (Ding and Gold, 2012), and working memory (Armstrong et al., 2009; Clark et al., 2012; Hasegawa et al., 2004).

It remains unclear how this heterogeneity in intrinsic timescales maps onto previously established functional and anatomical classifications of FEF neurons. Traditionally, FEF neurons have been categorized into visual, motor, visuomotor, and memory-delay neurons, based on their functional properties (Bruce and Goldberg, 1985; Thompson et al., 2005). Visual neurons likely have shorter intrinsic timescales consistent with their strong transient visual responses, whereas memory-delay neurons presumably have longer timescales, aligning with previous studies (Merrikhi et al., 2017; Wasmuht et al., 2018; Trepka et al., 2024). Furthermore, these long-*τ* neurons may also play a dominant role in attentional deployment, modulating visual cortical activity via their direct axonal projections (Burrows and Moore, 2009; Chen et al., 2020; Hü er et al., 2024; Soltani and Koch, 2010). Additionally, the laminar organization of the FEF may also contribute to the diversity in these timescales: granular-layer neurons receiving strong feedforward visual input have higher visual sensitivity and likely exhibit shorter timescales, whereas neurons in supra- and infra-granular layers, associated with feedback and integrative processing, may demonstrate longer timescales (Chen et al., 2018; Heinzle et al., 2007). Lastly, inhibitory and excitatory neurons may differ in their intrinsic timescales (Cavanagh et al., 2020). However, short-*τ* and long-*τ* neurons are unlikely to directly map to the inhibitory and excitatory neuron classes, as the ratio of short-*τ* to long-*τ* neurons is approximately 4:5, whereas the ratio of excitatory to inhibitory neurons in FEF is closer to 4:1 (Heinzle et al., 2007). Future research is needed to explore these relationships.

PPC inactivation had a selective impact on short-*τ* and long-*τ* neurons during different task epochs. Specifically, it had a substantially greater impact on the modulation of intrinsic timescales in short-*τ* neurons during the baseline period, and a stronger impact on attentional modulation in long-*τ* neurons during the task period. These differential effects suggest at least two distinct network motifs within the FEF, defined by separate contributions from short-*τ* and long-*τ* neurons (Fig. 6). Short-*τ* neurons exhibited strong transient visual responses and weak attentional modulation, implicating their primary role in relaying quickly changing visual information (Fig. 6A). These neurons are likely feedforward recipients, receiving direct projections from the PPC and other posterior visual regions such as those in the occipital lobes (Schall et al., 1995; Stanton et al., 1995), and have weaker recurrent connections with themselves and other PFC neurons. Consequently, PPC inactivation significantly altered their intrinsic dynamics but had a limited impact on their visual responses, possibly due to compensatory inputs from other visual regions. In contrast, long-*τ* neurons showed robust attentional modulation when presented with popout arrays (Fig. 6B). The input from the PPC and reciprocal connections between the PPC and FEF might represent or resolve stimulus competition and play an important role in salience computation (Buschman and Miller, 2007; Chen et al., 2020; Constantinidis and Steinmetz, 2005). This functional role might be driven by their recurrent connectivity within PFC that could promote sustained neural activity to maintain attentional modulation over time (Wang, 2020). Additionally, the relatively longer timescales of the PFC might dominate the dynamics of long-*τ* neurons in the FEF (Murray et al., 2014; Wasmuht et al., 2018), thus mitigating PPC’s influence on their intrinsic timescales. This would explain why PPC inactivation substantially impaired attentional modulation in these neurons, while having smaller effects on their intrinsic timescales. Future studies with large-scale recordings across both the PPC and FEF are essential to uncover the detailed circuit connections underlying these distinct neuronal populations.

**Fig. 6.**
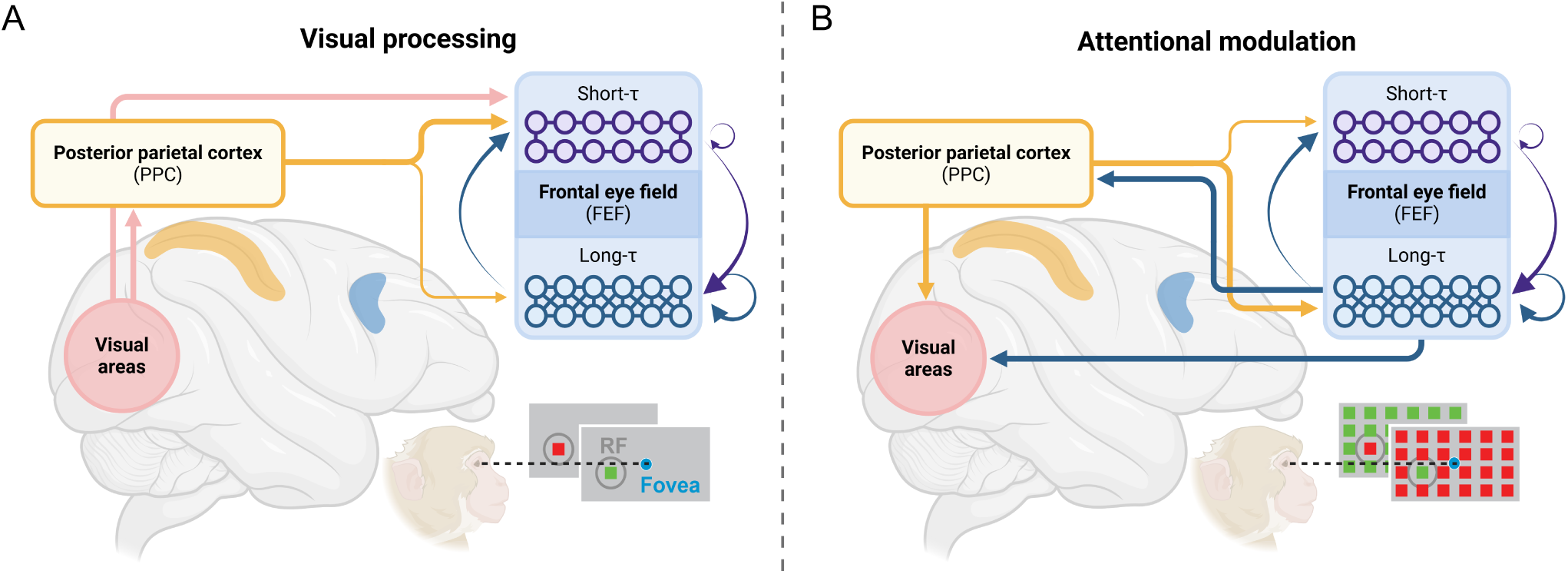
Proposed circuit mechanisms underlying visual processing and attentional modulation in the FEF. (A) Short-timescale (short-*τ*) neurons in the frontal eye field (FEF; blue) receive direct feedforward inputs from the posterior parietal cortex (PPC; orange) and early visual cortical areas (pink). These neurons possess relatively weaker recurrent connections and their intrinsic neural dynamics substantially depend on the PPC input. They predominantly relay sensory-driven visual information, as evidenced by their robust transient visual responses when presented with single visual stimuli. (B) Long-timescale (long-*τ*) neurons in the FEF are predominantly involved in attentional modulation. These neurons integrate information over larger visual space and temporal windows. Their role in salience computation in response to popout arrays significantly depends on the input from PPC. They likely maintain reciprocal connections with PPC, exhibit strong recurrent connectivity within prefrontal circuits, and project feedback signals to visual cortical areas. Consequently, they play a key role in sustaining attentional modulation and higher-order cognitive functions. Created in BioRender. Soyuhos, O. (2025) https://BioRender.com/g40r029.

Lastly, our findings provide direct empirical evidence demonstrating a causal role of long-range inter-area inputs in shaping intrinsic neural timescales within the FEF. Computational models have long suggested that intrinsic timescales emerge from interactions between local recurrent connectivity and long-range cortical projections, asserting that neither local nor long-range connectivity alone fully accounts for observed timescale variations (Chaudhuri et al., 2014, 2015; Demirtaş et al., 2019; Litwin-Kumar and Doiron, 2012; Wang, 2020). However, empirical tests of these model predictions have remained lacking. Here, the selective modulation of intrinsic timescales by PPC inactivation provides strong support for these theoretical predictions, emphasizing the importance of long-range cortical connectivity in shaping neuronal temporal dynamics. We propose two complementary mechanisms explaining the observed slowing of intrinsic timescales during PPC inactivation. First, parietal neurons exhibit faster neural dynamics compared to neurons in the frontal cortex (Chaudhuri et al., 2015; Murray et al., 2014; Wasmuht et al., 2018). The removal of PPC input might unmask inherently slower local dynamics within FEF neurons, particularly in short-*τ* neurons that have weaker recurrent structures. Second, PPC inactivation may enhance the engagement of local recurrent circuits within FEF or broader prefrontal cortical feedback loops, further prolonging neuronal integration times (Litwin-Kumar and Doiron, 2012). Future studies combining high-density neuronal recordings with targeted circuit manipulations will be crucial for testing these hypotheses, advancing our understanding of how intrinsic neural dynamics emerge from diverse network interactions and support cognitive functions.

## Methods

### Subject details

All experimental procedures followed the guidelines set by the National Institutes of Health Guide for the Care and Use of Laboratory Animals, the Society for Neuroscience Guidelines and Policies, and the Stanford University Animal Care and Use Committee. The experiments involved two healthy male rhesus macaques (Macaca mulatta): monkey Q (16 kg) and monkey J (17 kg). The number of animals used aligns with typical practices for neurophysiological studies involving non-human primates.

### Experimental procedures

The details of the surgical procedures and neurophysiological recordings have been previously reported by (Chen et al., 2020). In summary, two monkeys (Q, 16 kg; J, 17 kg) performed visual sensitivity and saliency-driven attention tasks under PPC-inactivation and control conditions while we monitored neural activity in the FEF (Fig. 1A). Each trial began with an initial central fixation period (1×1 dva) of 500 ms on a gray background (60 cd/m²), with the last 300 ms serving as the baseline period. Following this, the monkeys were presented with either a single stimulus or a popout stimulus for 500 ms while continuing to fixate at the center (Fig. 1B). In the single-stimulus trials, a red or green square stimulus (7×7 dva) was displayed at one of 24 locations on a 6×4 grid (75×45 dva). In the popout trials, a square stimulus was placed among an array of stimuli with contrasting colors (e.g., a red square among green stimuli or a green square among red stimuli). After the stimulus offset, the monkeys continued to fixate for an additional 500 ms to receive a reward (Fig. 1C).

### Neurophysiological recording of FEF neurons

The recording locations within the right FEF were identified by eliciting short-latency saccadic eye movements using biphasic current pulses (≤ 50 *µ*A; 250 Hz; 0.25 ms duration), in line with methods from previous studies (Bruce et al., 1985). Neural activity was recorded using 16- or 32-channel linear array electrodes (V and S-Probes, Plexon, Inc.), with contacts spaced 150 µm apart. These electrodes were inserted into the cortex via a hydraulic microdrive (Narishige International). Neural signals were referenced to a nearby stainless steel guide tube positioned near the electrode contacts. The signal used for spike detection was filtered with a 4-pole Bessel high-pass filter at 300 Hz and a 2-pole Bessel low-pass filter at 6000 Hz, and sampled at 40 kHz. Initial classification of extracellular waveforms into single neurons or multi-units was performed online via template matching, with offline sorting subsequently verified using Plexon systems. In total, we recorded spike timestamps from 400 neural units in the FEF across the two monkeys.

### Reversible inactivation of PPC

In each monkey, a pair of stainless steel cryoloops was implanted in the intraparietal sulcus (IPS) of the right hemisphere. These cryoloops were custom-designed to fit the contours of the IPS and fill the sulcus. The IPS was cooled by circulating chilled methanol through the loop tubing, with the temperature closely monitored and maintained within 1°C of the target value by adjusting the methanol flow rate. A stable loop temperature of approximately 5°C was reached within 5–10 minutes after the cooling process began, and the brain’s normal temperature was restored within about 2 minutes post-cooling. Temperatures around 5°C in the loop have been shown to effectively suppress neuronal activity in the underlying cortex (Lomber and Payne, 2000). During the experimental sessions, inactivation periods lasted 30 to 60 minutes each. Neurophysiological data were first collected during a control block, followed by an inactivation block across eight sessions. In six additional sessions, data collection followed two consecutive sets of control and inactivation phases (control–inactivation–control–inactivation), allowing the examination of potential block order effects. The success of PPC inactivation on behavior was evaluated in separate sessions using double-target choice tasks and free viewing of natural images (Chen et al., 2020).

### Computation of intrinsic neural timescales

We analyzed neural activity from 400 neural units in the FEF across two monkeys. Each unit’s activity was recorded as spike timestamps, which were binned into 10 ms intervals to calculate spike counts for each trial. To normalize the spike counts at each time bin across all trials, we subtracted the mean firing rate of each time bin from the spike count in that bin. This step helped isolate the intrinsic temporal dependencies of the neuron’s firing pattern, independent of its overall firing rate. We then performed autocorrelation analysis on the 300 ms baseline windows of each trial to compute the sum of products of spike count values against themselves at various lags for each neuron. The autocorrelation values for each lag were first averaged across trials for each neuron. Next, these averaged autocorrelation values were scaled by dividing them by the zero-lag value, standardizing the results for each neuron.

We then fitted a single exponential decay function to these averaged autocorrelation values and calculated the decay time constants for each neuron, representing their intrinsic neural timescales (Fig. 1D). Importantly, a subset of neurons displayed significantly low autocorrelation values at short time lags, suggesting the presence of refractory periods or negative adaptation, as reported in previous studies (Cavanagh et al., 2016; Murray et al., 2014; Wasmuht et al., 2018). Therefore, the fitting procedure was started at the time lag where the mean autocorrelation reached its maximum value (excluding the zeroth lag where autocorrelation equals one). For the curve fitting, we used the Trust Region Reflective algorithm. The model used for the fitting process is given by the equation:

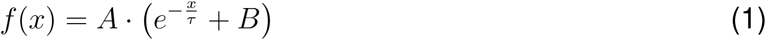

where *x* is the time lag, *τ* represents the intrinsic timescale, *A* is the amplitude of the decay, and *B* is included to adjust for baseline shifts, addressing the influence of intrinsic neural timescales that extend beyond the measurement window.

To establish an objective criterion for the goodness of fit, we calculated pseudo-*R*^2^ values for nonlinear fitting based on the proportion of the total variance explained by the exponential decay function. We found that 5% of units had *R*^2^ values below 0.3, indicating a poor fit. Additionally, 6.5% of units fell within the range of 0.3 to 0.5, representing a moderate fit, and 13% of units had values between 0.5 and 0.7, suggesting a good fit. The majority, 75.5% of units, demonstrated values ranging from 0.7 to 1.0, indicating an excellent model fit (Fig. S1). Neurons with *R*^2^ values below 0.3 were excluded from further analysis, resulting in a remaining sample of 380 neural units for the control condition and 372 neural units for the inactivation condition.

### Mode testing of intrinsic neural timescales

To investigate the distribution of intrinsic neural timescales in the FEF, we applied several statistical tests using the mode test function from the multimode R package (Ameijeiras-Alonso et al., 2021). Each test was chosen for its ability to highlight different aspects of distribution modality. We used the Hall and York Critical Bandwidth Test (Hall and York, 2001), which refines bandwidth selection in kernel density estimation to improve the detection of subtle multimodal patterns. The Fisher and Marron Cramer–von Mises Test (Fisher and Marron, 2001) was applied for its sensitivity in capturing subtle features like shoulder modes by assessing how well the empirical distribution function fits a hypothesized unimodal function using the Cramer–von Mises statistic. Additionally, we used the Excess Mass Test (Ameijeiras-Alonso et al., 2019), which employs the excess mass criterion for mode detection, offering a robust method particularly suited for multidimensional datasets with high noise levels. All tests provided complementary results (Table S1).

### Calculation of single and popout indices

We measured the receptive field of each neuron by analyzing their responses to a single stimulus presented at each of 24 locations. Locations where a neuron’s firing rate exceeded 95% of its maximum response were included within the neuron’s receptive field. In contrast, the four locations in the far-right column of the ipsilateral visual field were considered outside the receptive field. Following this, we computed visual indices for each neuron, reflecting the neural activity during transient (50–200 ms) and sustained (200–500 ms) periods for both single and popout stimulus trials. These indices were calculated based on the average differential spike rate when stimuli were presented within versus outside the neuron’s receptive field (In-RF vs. Out-RF; Fig. 1B), normalized by their sum.

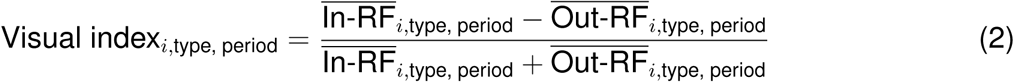

Here, *i* represents each neuron, ‘type’ refers to single or popout trials, and ‘period’ denotes transient or sustained periods. We computed four single and popout indices corresponding to transient and sustained visual responses as well as attentional modulation. The distributions of single and popout indices in the short-*τ* and long-*τ* neurons were compared using the Cramer–von Mises test (Fig. 3C).

### Analyzing visual onsets

We quantified the earliest time at which each neuron’s activity rose significantly above baseline (onset latency), separately for single and popout stimulus trials. Specifically, we computed the mean spike count across trials for each 10 ms bin. A baseline threshold was then defined by adding two standard deviations to the average spike count during the pre-stimulus window. Within the post-stimulus period, we then identified the first occurrence of at least two consecutive bins (20 ms) exceeding this threshold. The leading bin in this sequence was determined as the neuron’s onset bin. By applying this procedure independently to single and popout stimulus trials, we observed that 92% of short-*τ* neurons and 82% of long-*τ* neurons showed a detectable onset under single-stimulus trials. Additionally, 86% of short-*τ* neurons and 67% of long-*τ* neurons displayed a detectable onset during popout trials. These results provided onset latencies that characterized the timing of visually evoked responses across stimulus and experimental conditions (Fig. S6). Visual inspection of the neuronal activity traces confirmed the accuracy of these onset calculations.

### Multiple linear regression analyses

We conducted multiple regression analyses to assess the individual contributions of each variable while accounting for their interdependencies. In our model, the dependent variable was the intrinsic neural timescale, while the independent variables included the single indices, popout indices, and the baseline mean firing rate. The regression model was specified as:

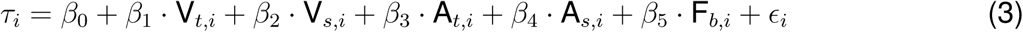

In this equation, *τ_i_*denotes the intrinsic neural timescale for neuron *i*; *V_t,i_* and *V_s,i_*are the transient and sustained single indices; *A_t,i_* and *A_s,i_* are the transient and sustained popout (stimulus-driven attention) indices; *F_b,i_* represents the baseline mean firing rate; and *ɛ_i_*accounts for unexplained variance.

To fit the linear model, we used robust regression, which minimizes the influence of outliers and is less sensitive to non-normality in the data. This approach ensures that the coefficients accurately reflect the effect of each predictor on the dependent variable without being disproportionately influenced by outliers. We utilized added-variable plots to illustrate the relationship between neural timescales and each predictor by plotting the controlled residuals against each independent variable (Fig. 3A). These plots provide a visual representation of the distinct contributions of each predictor to the model, with the slopes quantifying the effect of a one-unit change in each predictor on the timescales while holding the other variables constant.

We checked for multicollinearity among the independent variables in the multiple regression model using variance inflation factor (VIF) analysis. The VIF quantifies the degree to which the variance of a regression coefficient is inflated due to linear dependence on other predictors in the model. Generally, a VIF value greater than 5 or 10 suggests problematic multicollinearity (James et al., 2021), indicating that the affected predictors may be linearly dependent on others. In our analysis, we did not find significant multicollinearity (Fig. S2A).

Additionally, we examined the normality of the residuals in our model. A Q–Q plot was used to visually assess the distribution of residuals from our robust regression model against a theoretical normal distribution (Fig. S2B). Deviations from the line in this plot indicate departures from normality, suggesting that the residuals were not normally distributed. To address this, we used permutation testing to assess the significance of the regression coefficients. We employed a custom function to randomly shuffle the dependent variable (neural timescales) 10000 times while keeping the independent variables unchanged. This generated a distribution of regression coefficients under the null hypothesis of no association (Fig. 3A). P-values were calculated as the proportion of permuted coefficients whose absolute values equaled or exceeded that of the observed coefficient.

### Neuron-dropping decoding analyses

To assess whether short-*τ* and long-*τ* neurons differ in visual responses and attentional modulation at the population level, we conducted a neuron-dropping analysis (Lebedev, 2014). We tested the ability to decode spatial location (inside vs. outside the receptive field) during both transient and sustained stimulus presentations in both single-stimulus and popout trials. Neurons with fewer than 20 trials were excluded, resulting in 143 neurons in the short-*τ* group and 190 neurons in the long-*τ* group. To evaluate the decoding performance of varying neuron group sizes, we incrementally analyzed groups starting with three neurons and expanding up to 120 neurons. The maximum number of neurons was capped at 120 to avoid potential statistical validity issues that can arise when the subpopulation size approaches that of the fixed population (Lebedev, 2014). At each step, we randomly sampled the specified number of neurons 500 times to ensure robust performance estimates. For each decoding analysis (across 500 iterations at each step), we equalized the number of trials for In-RF and Out-RF conditions by randomly sampling an equal number of trials from each condition. Next, we conducted principal component analysis (PCA), retaining the top three principal components. A support vector machine (SVM) classifier was then used for decoding. Decoding performances exhibited saturation curves as the number of neurons increased, indicating diminishing returns in performance improvement with additional neurons.

To assess differences in growth rates between short-*τ* and long-*τ* neurons, we fit a saturation function defined as:

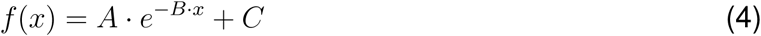

Here, *x* represents the number of neurons, ranging from 1 to 120; *A* is the initial amplitude; *B* is the saturation rate; and *C* is the asymptote value of the function. This approach allowed us to model and compare the dynamics of decoding performance as a function of population size between the two groups. To improve fitting accuracy, we averaged the results of every five iterations, resulting in 100 averaged data points for each group over the incremental steps up to 120 neurons. The saturation function was then fitted to these averaged data points for each condition, and the parameters *B* and *C* were compared between short-*τ* and long-*τ* neurons using the nonparametric Wilcoxon rank-sum test (Fig. S3).

Additionally, we analyzed the area under the curve (AUC) for decoding performance. The AUC was calculated for each group based on the same 100 averaged data points, measuring the area above the chance level. This provided an overall measure of decoding performance across all population sizes. To standardize the results, we normalized the AUC by dividing it by the maximum possible area above chance. A normalized score of 0 indicates overall performance at the chance level, whereas a score of 1 reflects perfect decoding accuracy across all population sizes. The AUC was computed using the trapezoidal rule, which finds the area between consecutive points to determine overall decoding performance. The resulting normalized AUC scores were then compared between short-*τ* and long-*τ* neurons using the nonparametric Wilcoxon rank-sum test (Fig. 3).

### Testing the effect of parietal inactivation

To investigate the effects of parietal inactivation on intrinsic neural timescales, visual responses, and attentional modulation in the FEF, we conducted a series of statistical analyses. First, we performed paired Wilcoxon signed-rank tests to determine whether parietal inactivation led to significant changes in FEF neural timescales, assessing the direction of change (increase or decrease) across both monkeys and within each individual monkey. The rank biserial correlation effect size was calculated to quantify the magnitude of the observed changes (Fig. 4B). Next, we analyzed whether the change (Δ: inactivation – control) in neural timescales differed between short-*τ* and long-*τ* neurons. We first tested against zero using a one-sample Wilcoxon signed-rank test to determine whether the change within each group was significant, followed by unpaired Wilcoxon rank-sum tests to compare the magnitude of change between the two groups (Fig. 4E).

We also examined whether the relationships between neural timescales and visual/attentional responses observed in the control condition persisted or were altered during inactivation. A multiple regression analysis was conducted using values obtained after inactivation, with intrinsic neural timescales as the dependent variable and visual indices along with baseline firing rate as independent variables (Fig. 5A). Additionally, changes in the distribution of single and popout indices were compared between short-*τ* and long-*τ* neurons using the Cramer–von Mises test (Fig. 5). Furthermore, we performed a neuron-dropping analysis (Lebedev, 2014) under PPC inactivation to evaluate changes in single- and popout-decoding performance at the population level (Fig. 5C–D). The function parameters (saturation rate and asymptote value) were compared between short-*τ* and long-*τ* neurons in both conditions (Fig. S5). Lastly, we compared changes in the normalized AUC scores between the two groups following PPC inactivation (Fig. 5E).

### Data and code availability

The dataset and code used in this study will be made publicly available on Figshare and Code Ocean following acceptance of this manuscript.

## Acknowledgments

We thank Dr. Tirin Moore for his contributions to the initial experimental design, securing funding for the project, and for providing constructive feedback. This work was supported by the National Science Foundation under grants 2152260 (NSF NRT NeuralStorm) (to O.S.) and 2207895 (to R.C.), National Institutes of Health grants EY014924 (to X.C.) and EY029759 (to T.M.), the Brain and Behavior Research Foundation (to X.C.), the Alfred P. Sloan Foundation under Grant No. FG-2021-16304 (to R.C.), and the UC Davis Large Interdisciplinary Applications in Neuroscience (LIAN) (to R.C. and X.C.).

## Author information

### Contributions

Study conceptualization: X.C., O.S., R.C. and T.M.; Experimental design: X.C., M.Z., and T.M.; Performing experiments: X.C. and M.Z.; Formal analysis: O.S.; Visualization: O.S.; Writing – original draft: O.S. and X.C.; Writing – review & editing: O.S., X.C, R.C. and T.M.; Funding acquisition: X.C, T.M., O.S. and R.C.

### Corresponding author

Correspondence to Xiaomo Chen

## References

Ameijeiras-Alonso, J., Crujeiras, R. M., and Rodriguez-Casal, A. (2021). multimode: An R Package for Mode Assessment. Journal of Statistical Software, 97:1–32.

Ameijeiras-Alonso, J., Crujeiras, R. M., and Rodríguez-Casal, A. (2019). Mode testing, critical bandwidth and excess mass. TEST, 28(3):900–919.

Anderson, J. C., Kennedy, H., and Martin, K. A. C. (2011). Pathways of Attention: Synaptic Relationships of Frontal Eye Field to V4, Lateral Intraparietal Cortex, and Area 46 in Macaque Monkey. Journal of Neuroscience, 31(30):10872–10881.

Armstrong, K. M., Chang, M. H., and Moore, T. (2009). Selection and Maintenance of Spatial Information by Frontal Eye Field Neurons. Journal of Neuroscience, 29(50):15621–15629.

Boshra, R. and Kastner, S. (2022). Attention control in the primate brain. Current Opinion in Neurobiology, 76:102605.

Bruce, C. J. and Goldberg, M. E. (1985). Primate frontal eye fields. I. Single neurons discharging before saccades. Journal of Neurophysiology, 53(3):603–635.

Bruce, C. J., Goldberg, M. E., Bushnell, M. C., and Stanton, G. B. (1985). Primate frontal eye fields. II. Physiological and anatomical correlates of electrically evoked eye movements. Journal of Neurophysiology, 54(3):714–734.

Burrows, B. E. and Moore, T. (2009). Influence and Limitations of Popout in the Selection of Salient Visual Stimuli by Area V4 Neurons. Journal of Neuroscience, 29(48):15169–15177. Publisher: Society for Neuroscience Section: Articles.

Buschman, T. J. and Miller, E. K. (2007). Top-Down Versus Bottom-Up Control of Attention in the Prefrontal and Posterior Parietal Cortices. Science, 315(5820):1860–1862.

Cavanagh, S. E., Hunt, L. T., and Kennerley, S. W. (2020). A Diversity of Intrinsic Timescales Underlie Neural Computations. Frontiers in Neural Circuits, 14.

Cavanagh, S. E., Wallis, J. D., Kennerley, S. W., and Hunt, L. T. (2016). Autocorrelation structure at rest predicts value correlates of single neurons during reward-guided choice. eLife, 5:e18937.

Chaudhuri, R., Bernacchia, A., and Wang, X.-J. (2014). A diversity of localized timescales in network activity. eLife, 3:e01239.

Chaudhuri, R., Knoblauch, K., Gariel, M.-A., Kennedy, H., and Wang, X.-J. (2015). A Large-Scale Circuit Mechanism for Hierarchical Dynamical Processing in the Primate Cortex. Neuron, 88(2):419–431.

Chen, X., Zirnsak, M., and Moore, T. (2018). Dissonant Representations of Visual Space in Prefrontal Cortex during Eye Movements. Cell Reports, 22(8):2039–2052.

Chen, X., Zirnsak, M., Vega, G. M., Govil, E., Lomber, S. G., and Moore, T. (2020). Parietal Cortex Regulates Visual Salience and Salience-Driven Behavior. Neuron, 106(1):177–187.e4.

Clark, K. L., Noudoost, B., and Moore, T. (2012). Persistent Spatial Information in the Frontal Eye Field during Object-Based Short-Term Memory. Journal of Neuroscience, 32(32):10907– 10914.

Constantinidis, C. and Steinmetz, M. A. (2005). Posterior Parietal Cortex Automatically Encodes the Location of Salient Stimuli. Journal of Neuroscience, 25(1):233–238.

Demirtaş, M., Burt, J. B., Helmer, M., Ji, J. L., Adkinson, B. D., Glasser, M. F., Van Essen, D. C., Sotiropoulos, S. N., Anticevic, A., and Murray, J. D. (2019). Hierarchical Heterogeneity across Human Cortex Shapes Large-Scale Neural Dynamics. Neuron, 101(6):1181–1194.e13.

Dias, E. C. and Segraves, M. A. (1999). Muscimol-Induced Inactivation of Monkey Frontal Eye Field: Effects on Visually and Memory-Guided Saccades. Journal of Neurophysiology, 81(5):2191–2214.

Ding, L. and Gold, J. I. (2012). Neural correlates of perceptual decision making before, during, and after decision commitment in monkey frontal eye field. Cerebral Cortex (New York, N.Y.: 1991), 22(5):1052–1067.

Elston, G., Benavides-Piccione, R., Elston, A., Manger, P., and Defelipe, J. (2011). Pyramidal Cells in Prefrontal Cortex of Primates: Marked Differences in Neuronal Structure Among Species. Frontiers in Neuroanatomy, 5.

Elston, G. N. (2000). Pyramidal Cells of the Frontal Lobe: All the More Spinous to Think With. Journal of Neuroscience, 20(18):RC95–RC95.

Elston, G. N. (2003). Cortex, cognition and the cell: new insights into the pyramidal neuron and prefrontal function. Cerebral Cortex (New York, N.Y.: 1991), 13(11):1124–1138.

Fallon, J., Ward, P. G. D., Parkes, L., Oldham, S., Arnatkevičiūtė, A., Fornito, A., and Fulcher, B. D. (2020). Timescales of spontaneous fMRI fluctuations relate to structural connectivity in the brain. Network Neuroscience, 4(3):788–806.

Fiebelkorn, I. C. and Kastner, S. (2020). Functional Specialization in the Attention Network. Annual Review of Psychology, 71(Volume 71, 2020):221–249.

Fiebelkorn, I. C., Pinsk, M. A., and Kastner, S. (2018). A Dynamic Interplay within the Frontoparietal Network Underlies Rhythmic Spatial Attention. Neuron, 99(4):842–853.e8.

Fisher, N. I. and Marron, J. S. (2001). Mode testing via the excess mass estimate. Biometrika, 88(2):499–517.

Gao, R., van den Brink, R. L., Pfeffer, T., and Voytek, B. (2020). Neuronal timescales are functionally dynamic and shaped by cortical microarchitecture. eLife, 9:e61277.

Golesorkhi, M., Gomez-Pilar, J., Zilio, F., Berberian, N., Wolff, A., Yagoub, M. C. E., and Northoff, G. (2021). The brain and its time: intrinsic neural timescales are key for input processing. Communications Biology, 4(1):1–16.

Hall, P. and York, M. (2001). On the Calibration of Silverman’s Test for Multimodality. Statistica Sinica, 11(2):515–536.

Hart, E. and Huk, A. C. (2020). Recurrent circuit dynamics underlie persistent activity in the macaque frontoparietal network. eLife, 9:e52460.

Hasegawa, R. P., Peterson, B. W., and Goldberg, M. E. (2004). Prefrontal Neurons Coding Suppression of Specific Saccades. Neuron, 43(3):415–425.

Hasson, U., Chen, J., and Honey, C. J. (2015). Hierarchical process memory: memory as an integral component of information processing. Trends in cognitive sciences, 19(6):304–313.

Heinzle, J., Hepp, K., and Martin, K. A. C. (2007). A Microcircuit Model of the Frontal Eye Fields. Journal of Neuroscience, 27(35):9341–9353.

Hüer, J., Saxena, P., and Treue, S. (2024). Pathway-selective optogenetics reveals the functional anatomy of top–down attentional modulation in the macaque visual cortex. Proceedings of the National Academy of Sciences, 121(3):e2304511121.

James, G., Witten, D., Hastie, T., and Tibshirani, R. (2021). Linear Regression. In James, G., Witten, D., Hastie, T., and Tibshirani, R., editors, An Introduction to Statistical Learning: with Applications in R, pages 59–128. Springer US, New York, NY.

Lebedev, M. A. (2014). How to read neuron-dropping curves? Frontiers in Systems Neuroscience, 8:102.

Litwin-Kumar, A. and Doiron, B. (2012). Slow dynamics and high variability in balanced cortical networks with clustered connections. Nature Neuroscience, 15(11):1498–1505.

Lomber, S. G. and Payne, B. R. (2000). Translaminar differentiation of visually guided behaviors revealed by restricted cerebral cooling deactivation. Cerebral Cortex (New York, N.Y.: 1991), 10(11):1066–1077.

Lynch, J. C. and Tian, J.-R. (2006). Cortico-cortical networks and cortico-subcortical loops for the higher control of eye movements. Progress in Brain Research, 151:461–501.

Merrikhi, Y., Clark, K., Albarran, E., Parsa, M., Zirnsak, M., Moore, T., and Noudoost, B. (2017). Spatial working memory alters the efficacy of input to visual cortex. Nature Communications, 8(1):15041.

Moore, T. and Fallah, M. (2001). Control of eye movements and spatial attention. Proceedings of the National Academy of Sciences of the United States of America, 98(3):1273–1276.

Moore, T. and Fallah, M. (2004). Microstimulation of the frontal eye field and its effects on covert spatial attention. Journal of Neurophysiology, 91(1):152–162.

Moore, T. and Zirnsak, M. (2017). Neural Mechanisms of Selective Visual Attention. Annual Review of Psychology, 68(Volume 68, 2017):47–72.

Murray, J. D., Bernacchia, A., Freedman, D. J., Romo, R., Wallis, J. D., Cai, X., Padoa-Schioppa, C., Pasternak, T., Seo, H., Lee, D., and Wang, X.-J. (2014). A hierarchy of intrinsic timescales across primate cortex. Nature Neuroscience, 17(12):1661–1663.

Murray, J. D., Jaramillo, J., and Wang, X.-J. (2017). Working Memory and Decision-Making in a Frontoparietal Circuit Model. Journal of Neuroscience, 37(50):12167–12186.

Pettine, W. W., Steinmetz, N. A., and Moore, T. (2019). Laminar segregation of sensory coding and behavioral readout in macaque V4. Proceedings of the National Academy of Sciences, 116(29):14749–14754.

Raut, R. V., Snyder, A. Z., and Raichle, M. E. (2020). Hierarchical dynamics as a macroscopic organizing principle of the human brain. Proceedings of the National Academy of Sciences, 117(34):20890–20897.

Schall, J. D., Morel, A., King, D. J., and Bullier, J. (1995). Topography of visual cortex connections with frontal eye field in macaque: convergence and segregation of processing streams. The Journal of Neuroscience: The Official Journal of the Society for Neuroscience, 15(6):4464– 4487.

Soltani, A. and Koch, C. (2010). Visual Saliency Computations: Mechanisms, Constraints, and the Effect of Feedback. Journal of Neuroscience, 30(38):12831–12843.

Soltani, A., Murray, J. D., Seo, H., and Lee, D. (2021). Timescales of cognition in the brain. Current Opinion in Behavioral Sciences, 41:30–37.

Spitmaan, M., Seo, H., Lee, D., and Soltani, A. (2020). Multiple timescales of neural dynamics and integration of task-relevant signals across cortex. Proceedings of the National Academy of Sciences, 117(36):22522–22531.

Stanton, G. B., Bruce, C. J., and Goldberg, M. E. (1995). Topography of projections to posterior cortical areas from the macaque frontal eye fields. Journal of Comparative Neurology, 353(2):291–305.

Szczepanski, S. M., Konen, C. S., and Kastner, S. (2010). Mechanisms of Spatial Attention Control in Frontal and Parietal Cortex. Journal of Neuroscience, 30(1):148–160.

Thompson, K. G., Biscoe, K. L., and Sato, T. R. (2005). Neuronal Basis of Covert Spatial Attention in the Frontal Eye Field. Journal of Neuroscience, 25(41):9479–9487.

Trepka, E., Spitmaan, M., Qi, X.-L., Constantinidis, C., and Soltani, A. (2024). Training-Dependent Gradients of Timescales of Neural Dynamics in the Primate Prefrontal Cortex and Their Contributions to Working Memory. Journal of Neuroscience, 44(2).

Wang, X.-J. (2008). Decision Making in Recurrent Neuronal Circuits. Neuron, 60(2):215–234.

Wang, X.-J. (2020). Macroscopic gradients of synaptic excitation and inhibition in the neocortex. Nature Reviews. Neuroscience, 21(3):169–178.

Wasmuht, D. F., Spaak, E., Buschman, T. J., Miller, E. K., and Stokes, M. G. (2018). Intrinsic neuronal dynamics predict distinct functional roles during working memory. Nature Communications, 9(1):3499.

Wessberg, J., Stambaugh, C. R., Kralik, J. D., Beck, P. D., Laubach, M., Chapin, J. K., Kim, J., Biggs, S. J., Srinivasan, M. A., and Nicolelis, M. A. L. (2000). Real-time prediction of hand trajectory by ensembles of cortical neurons in primates. Nature, 408(6810):361–365.

Wolff, A., Berberian, N., Golesorkhi, M., Gomez-Pilar, J., Zilio, F., and Northoff, G. (2022). Intrinsic neural timescales: temporal integration and segregation. Trends in Cognitive Sciences, 26(2):159–173.

Wong, K.-F. and Wang, X.-J. (2006). A recurrent network mechanism of time integration in perceptual decisions. The Journal of Neuroscience: The Official Journal of the Society for Neuroscience, 26(4):1314–1328.

Xia, R., Chen, X., Engel, T. A., and Moore, T. (2024). Common and distinct neural mechanisms of attention. Trends in Cognitive Sciences, 28(6):554–567.

Zeraati, R., Shi, Y.-L., Steinmetz, N. A., Gieselmann, M. A., Thiele, A., Moore, T., Levina, A., and Engel, T. A. (2023). Intrinsic timescales in the visual cortex change with selective attention and reflect spatial connectivity. Nature Communications, 14(1):1858.

