## Supplementary information for "Selective control of prefrontal neural timescales by parietal cortex"

#### Figures

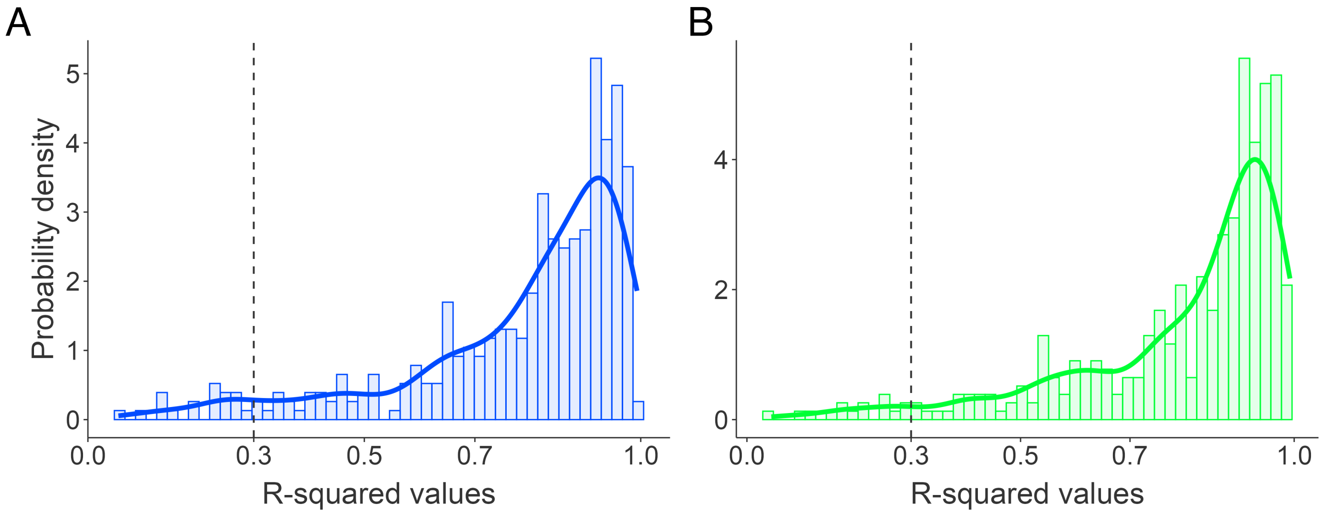


**Fig. S1 | Distribution of R-squared values from exponential decay fittings, related to Fig. 1C.**

The histograms show the $R^2$ values obtained from fitting an exponential decay function to the autocorrelation values of each neuron for control (blue) and PPC inactivation (green) conditions.

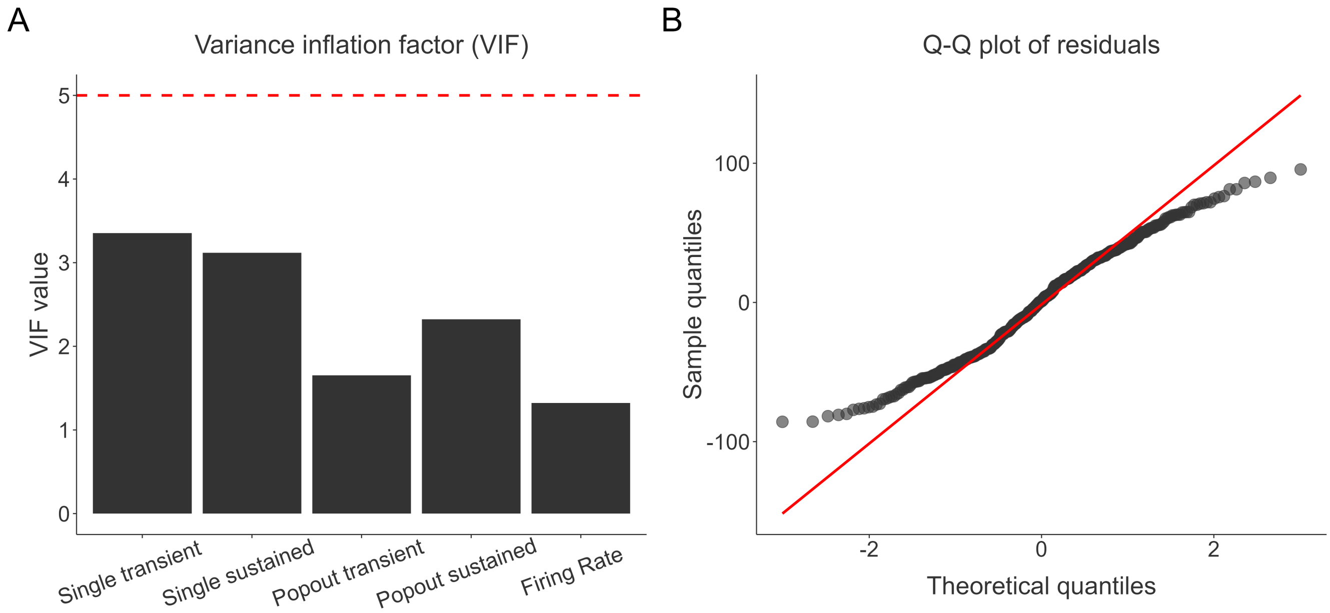


#### Fig. S2 | Diagnostics for the multiple regression model, related to Fig. 3A.

(A) Variance inflation factor (VIF) plot showing the VIF for each predictor. The red line indicates potential multicollinearity for VIF values exceeding 5. (B) Quantile-Quantile (Q-Q) plot examining the normality of model residuals. Departures from the red reference line suggest deviations from a normal distribution.

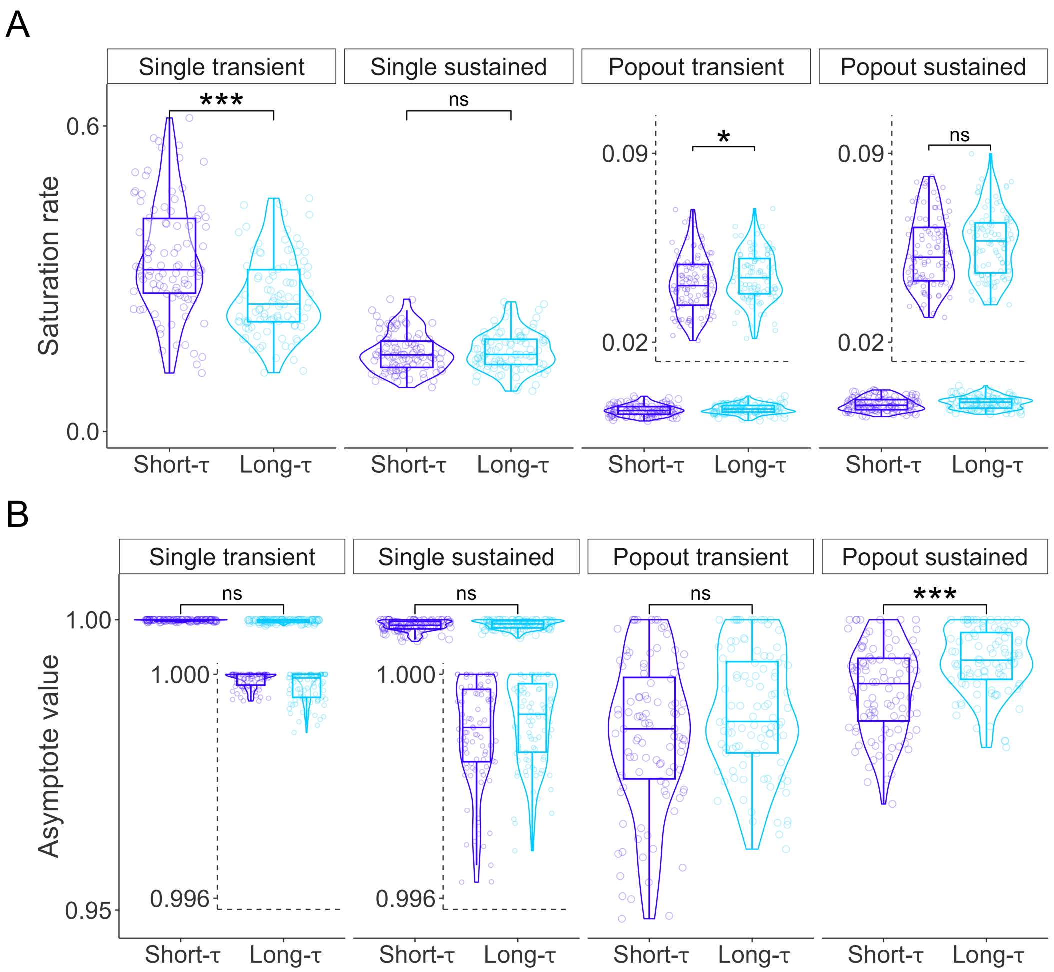


**Fig. S3 | Comparison of saturation rates and asymptotic values, related to Fig. 3D-E**

(A) Boxplots comparing saturation rates between short-timescale (short-$\tau$) and long-timescale (long-$\tau$) neurons for single- and popout-decoding performance, derived from neuron-dropping curves under the control condition. (B) Same as (A), but for asymptotic values. (*, $p < 0.05$; **, $p < 0.01$; ***, $p < 0.001$; ns = not significant).

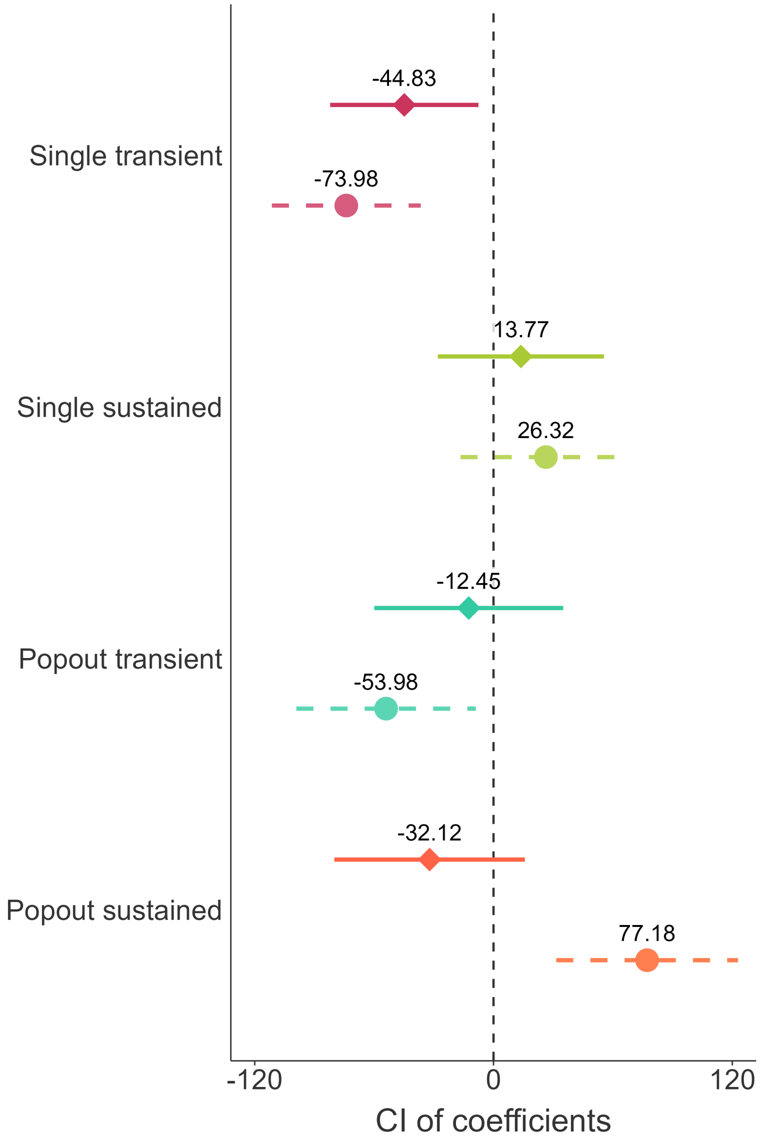


**Fig. S4 | Changes in confidence intervals for multiple regression models, related to Fig. 5A.**

The figure compares confidence intervals (CIs) of regression coefficients between control (dashed lines, circles) and inactivation (solid lines, diamonds) conditions. The vertical dashed line at zero indicates significance; predictors with CIs not crossing zero have a significant relationship with intrinsic neural timescales. Circles and diamonds represent the observed coefficients for control and inactivation, respectively.

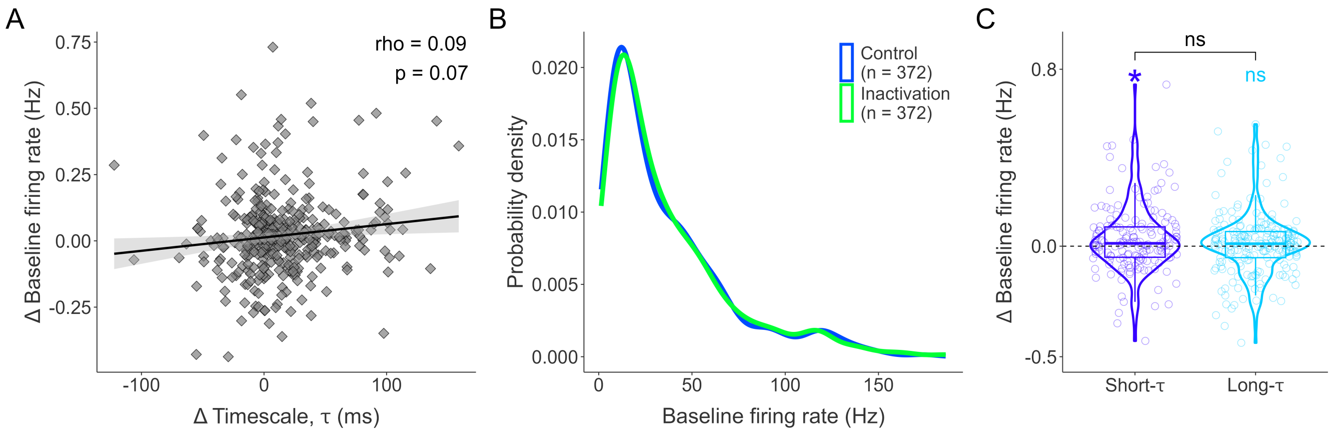


**Fig. S5 | Baseline firing rate and its relationship with intrinsic neural timescales following PPC inactivation, related to Fig. 4.**

(A) Correlation between change ($\Delta$) in intrinsic timescales ($\tau$) and baseline firing rates across neurons. (B) Probability density plots showing the distributions of baseline firing rate during control (blue) and inactivation (green) conditions. (C) Violin plots showing change in baseline firing rates for short-timescale (short-$\tau$) and long-timescale (long-$\tau$) neurons following PPC inactivation. (*, $p < 0.05$; **, $p < 0.01$; ***, $p < 0.001$; ns = not significant).

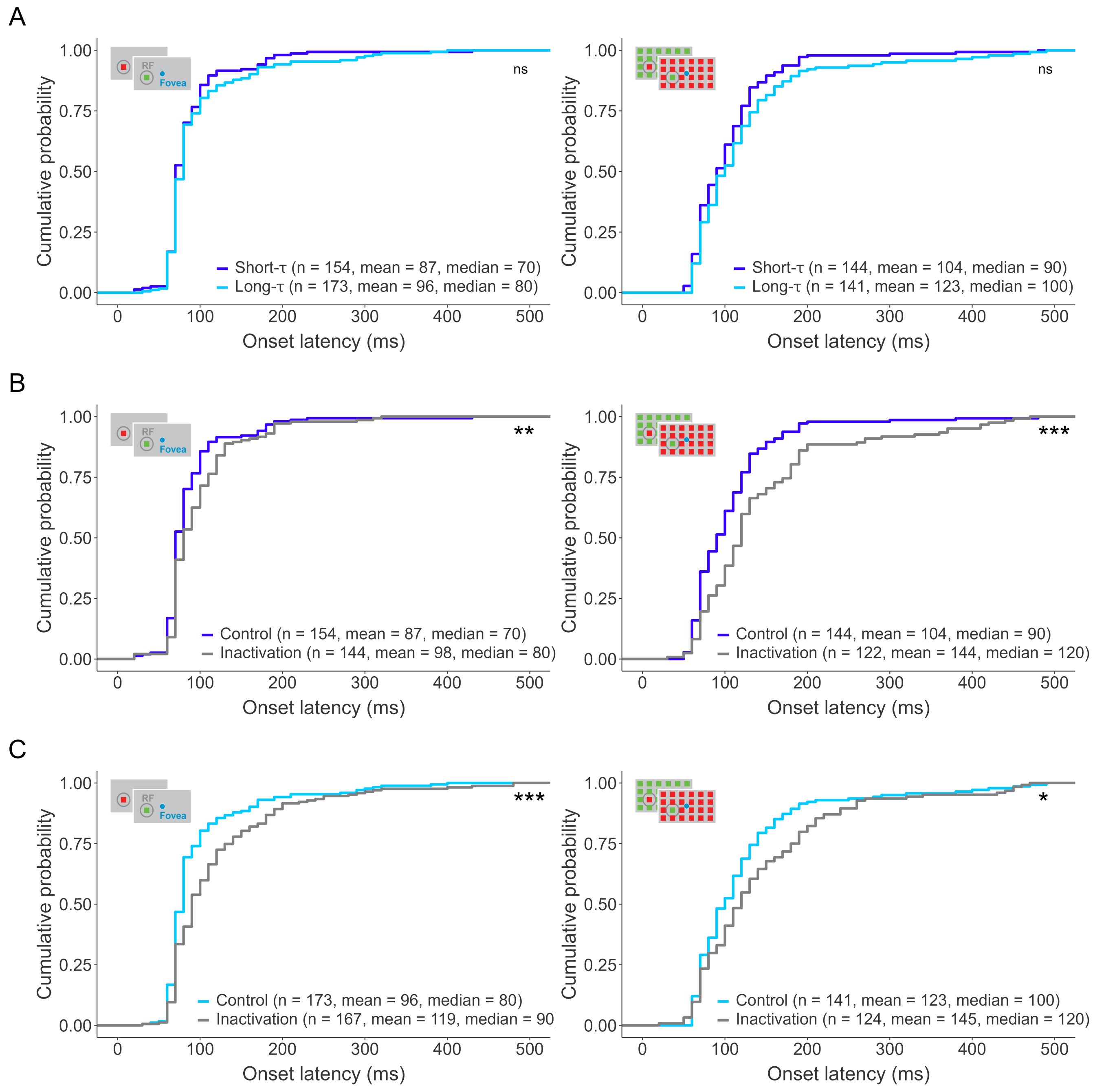


**Fig. S6 | Cumulative distributions of visual onset latencies, related to Fig. 3B.**

(A) Cumulative distributions of onset latencies (ms) for short-$\tau$ (purple) and long-$\tau$ (blue) neurons during single-stimulus (left) and popout (right) trials. (B) Cumulative distributions for short-$\tau$ neurons, comparing control (purple) vs. PPC inactivation (gray) conditions during single-stimulus (left) and popout (right) trials. (C) As in (B), but for long-$\tau$ neurons. (*, $p < 0.05$; **, $p < 0.01$; ***, $p < 0.001$; ns = not significant).

### Tables

| Method | Reference | Monkey(s) | Test Statistic | P-value |
|---|---|---|---|---|
| Ameijeiras-Alonso et al. Excess Mass Test | Ameijeiras-Alonso et al., 2019 | Both monkeys | 0.07 | $p < 0.001$ |
| | | Monkey J | 0.07 | $p < 0.05$ |
| | | Monkey Q | 0.09 | $p < 0.001$ |
| Fisher and Marron Cramer-von Mises Test | Fisher & Marron, 2001 | Both monkeys | 0.48 | $p < 0.001$ |
| | | Monkey J | 0.21 | $p < 0.05$ |
| | | Monkey Q | 0.42 | $p < 0.001$ |
| Hall and York Critical Bandwidth Test | Hall & York, 2001 | Both monkeys | 26.27 | $p < 0.001$ |
| | | Monkey J | 24.15 | $p < 0.05$ |
| | | Monkey Q | 27.97 | $p < 0.001$ |

#### Table S1 | Modality test results for intrinsic neural timescales, related to Fig. 2.

The table reports the results of statistical tests assessing the modality of intrinsic neural timescale distributions in the frontal eye field (FEF). Test statistics and corresponding P-values are provided for both monkeys combined, as well as separately for Monkey J and Monkey Q.
